# An examination of the divergent spatiotemporal signaling of GLP-1R *versus* GIPR in pancreatic beta cells

**DOI:** 10.1101/2022.08.17.504231

**Authors:** Yusman Manchanda, Stavroula Bitsi, Shiqian Chen, Johannes Broichhagen, Jorge Bernardino de la Serna, Ben Jones, Alejandra Tomas

## Abstract

The incretin receptors, glucagon-like peptide-1 receptor (GLP-1R) and glucose-dependent insulinotropic polypeptide receptor (GIPR), are class B GPCRs and prime therapeutic targets for the treatment of type 2 diabetes (T2D) and obesity. They are expressed in pancreatic beta cells where they potentiate insulin release in response to food intake. Despite GIP being the main incretin in healthy individuals, GLP-1R has been favoured *versus* GIPR as a therapeutic target due to GIPR responses being blunted in T2D patients and the conflicting effects of GIPR agonists and antagonists in improving glucose tolerance and preventing weight gain. There is, however, a recently renewed interest in GIPR biology following the realisation that GIPR responses can be restored after an initial period of blood glucose normalization and the recent development of dual GLP-1R-GIPR agonists with superior capacity for the control of blood glucose levels and weight. The importance of GLP-1R trafficking and subcellular signaling in the control of receptor outputs is well established, but little is known about the pattern of spatiotemporal signaling from the GIPR in beta cells. Here we have directly compared the main trafficking and signaling characteristics of both receptors in pancreatic beta cells, finding striking differences in their propensities for internalization, recycling, and degradation, as well as plasma membrane *versus* endosomal activity, with potential implications for receptor-specific control of beta cell function.

## Introduction

Incretin receptors, comprising the glucagon-like peptide-1 receptor (GLP-1R) and the glucose-dependent insulinotropic polypeptide receptor (GIPR), are key components of the glucoregulatory system due to their capacity to prevent postprandial hyperglycemia by amplifying insulin secretion from pancreatic beta cells in a glucose-dependent manner [1]. The GLP-1R has successfully been exploited for the treatment of type 2 diabetes (T2D), with a number of pharmacological GLP-1R agonists currently in use clinically or undergoing clinical trials [2]. On the other hand, the GIPR has not until recently been intensively pursued as a T2D treatment target, primarily due to GIP responses being blunted in T2D patients [3] and the perception that GIPR activation leads to weight gain, as inferred from the observation that GIPR knockout (KO) mice are protected against the effects of an obesogenic diet [4]. As a result, antagonizing rather than activating the GIPR has been suggested as a potential therapeutic intervention for diabetes and obesity. However, recent data from preclinical and clinical studies appear to contradict these assumptions regarding the role of GIPR in diabetes, as GIPR agonists have been shown to improve glucose tolerance and reduce body weight in T2D patients [5], and dual GLP-1R – GIPR targeting peptides, such as the recently developed tirzepatide, have demonstrated enhanced efficacy compared to currently approved GLP-1R agonist monotherapies [6].

Previous studies from our group and others have demonstrated that changes in the spatiotemporal regulation of signaling play a crucial role in determining the metabolic outcomes of GLP-1R activation [7], and underlie the enhanced therapeutic effects of partial or biased GLP-1R agonists [8]. However, neither the GIPR trafficking nor the spatiotemporal regulation of signaling by active GIPRs have been well characterized, despite the potential importance of these processes in the paradoxical responses obtained with both GIPR agonists and antagonists. Moreover, GIPR behaviors have seldom been compared directly to the GLP-1R in pancreatic beta cell systems, the primary cell type in which these related receptors co-exist and exert many of their metabolic effects.

In the present study, we present a dataset describing the effects of GLP-1R *versus* GIPR activation on target downregulation and compartmentalization of intracellular signaling responses specifically in pancreatic beta cells, unveiling striking differences between the trafficking and signaling signatures from the two incretin receptors within their native environment.

## Materials and Methods

### Peptides

Native sequence peptides including GLP-1(7-36)NH_2_ (referred to as GLP-1) and GIP(1-42) (referred to as GIP), and their fluorescein isothiocyanate (FITC) and tetramethylrhodamine (TMR) conjugates were obtained from Wuxi Apptec at >90% purity. Full amino acid sequences are shown in Supplementary Figure 1E (FITC conjugates) and Supplementary Figure 3A (TMR conjugates).

### Cell culture

Parental INS-1 832/3 cells (a gift from Prof Christopher Newgard, Duke University, USA), INS-1 832/3 cells with endogenous GLP-1R or GIPR deleted by CRISPR/Cas9 [9] (a gift from Dr Jacqueline Naylor, MedImmune), and corresponding INS-1 832/3 cells stably expressing SNAP-GLP-1R or SNAP-GIPR (generated by transfecting GLP-1R KO or GIPR KO cells with SNAP-GLP-1R or SNAP-GIPR constructs (Cisbio), respectively, followed by selection with 1 mg/mL G418, FACS sorting of SNAP-receptor expressing cells and maintenance in 0.5 mg/mL G418) were cultured in RPMI-1640 with 11 mM D-glucose, supplemented with 10% FBS, 10 mM HEPES, 1 mM sodium pyruvate, 50 μM β-mercaptoethanol, and 1% penicillin/streptomycin in a 37ºC / 5% CO_2_ incubator.

### DERET internalization assays

The assay was performed as previously described [10]. INS-1 832/3 SNAP-GLP-1R or SNAP-GIPR cells were labelled in suspension with Lumi4-Tb (40 nM) for 30 min in complete media. After washing, cells were resuspended in HBSS containing 24 µM fluorescein and dispensed into 96-well white plates. A baseline read was serially recorded over 5 min using a Flexstation 3 instrument at 37°C in TR-FRET mode using the following settings: λ_ex_ 340 nm, λ_em_ 520 and 620 nm, auto-cut-off, delay 400 µs, integration time 1500 µs. Ligands were then added, after which signal was repeatedly recorded for 30 min. Fluorescence signals were expressed ratiometrically after first subtracting signal from wells containing 24 µM fluorescein but no cells. Internalization was quantified as area under the curve (AUC) relative to individual well baseline.

### High-content microscopy assays for receptor internalization and recycling

The assay was performed as previously described [10]. INS-1 832/3 SNAP-GLP-1R or SNAP-GIPR cells were seeded into poly-D-lysine-coated, black 96-well plates. On the day of the assay, labelling was performed with BG-S-S-649 (1 µM), a surface-labelling SNAP-tag probe that can be released on application of reducing agents such as Mesna. After washing, treatments were applied for 30 min at 37°C in complete medium. Ligand was removed and cells washed with cold HBSS and placed on ice for subsequent steps. Mesna (100 mM in alkaline TNE buffer, pH 8.6) or alkaline TNE buffer without Mesna was applied for 5 min, and then washed with HBSS. Cells were imaged by widefield microscopy, with both epifluorescence and transmitted phase contrast images acquired. On imaging completion, HBSS was removed and replaced with fresh complete medium, and receptor was allowed to recycle for 60 min at 37°C, followed by a second Mesna application to remove any receptor that had recycled to the plasma membrane, with the plate re-imaged as above. Internalized receptor at each time-point was determined from cell-containing regions as determined from the phase contrast image using PHANTAST [11], and used to determine internalization and recycling parameters as previously described [10].

### NanoBiT complementation and NanoBRET assays

#### Mini-G protein/β-arrestin-2 recruitment NanoBiT assays

Here the SmBiT was cloned in frame at the C-terminus of the GLP-1R and the GIPR by substitution of the Tango sequence on FLAG-tagged GLP-1R-Tango or GIPR-Tango (a gift from Prof Bryan Roth, University of North Carolina, USA; Addgene plasmids #66291 and #66294), respectively. Mini-Gs, mini-Gq, and mini-Gi plasmids, tagged at the N-terminus with LgBiT, were a gift from Prof Nevin Lambert, Medical College of Georgia, USA. For β-arrestin 2 recruitment assays, β-arrestin 2 fused at the N-terminus to LgBiT (LgBiT-β-arrestin 2; Promega, plasmid no. CS1603B118) was chosen as it has previously been used successfully with other class B GPCRs. INS-1 832/3 GLP-1R KO and GIPR KO cells were seeded in 12-well plates and co-transfected with 0.5 μg each of GLP-1R-SmBiT or GIPR-SmBiT and either LgBiT-mini-Gs, -mini-Gq, -mini-Gi or -β-arrestin 2.

#### KRAS/Rab5 bystander NanoBRET assays

GLP-1R-NanoLuc was generated *in house* by PCR cloning of the NanoLuciferase sequence from pcDNA3.1-ccdB-NanoLuc (a gift from Prof Mikko Taipale; Addgene plasmid # 87067) onto the C-terminus end of the SNAP-GLP-1R vector (CisBio), followed by site-directed mutagenesis of the GLP-1R stop codon. GIPR-NanoLuc was subsequently cloned *in house* by exchanging the GLP-1R for the GIPR in the GLP-1R-NanoLuc construct. KRAS- and Rab5-Venus plasmids were a gift from Prof Kevin Pfleger, University of Western Australia. INS-1 832/3 GLP-1R KO and GIPR KO cells were seeded in 12-well plates and co-transfected with 0.2 µg KRAS-Venus and 0.1 µg GLP-1R- or GIPR-NanoLuc, respectively, or 0.5 µg Rab5-Venus and 0.1 µg GLP-1R- or GIPR-NanoLuc, respectively.

#### KRAS/Rab5 bystander mini-Gs NanoBRET assays

Mini-Gs-NanoLuc was a gift from Prof Nevin Lambert, Augusta University, USA. INS-1 832/3 GLP-1R KO or GIPR KO cells were seeded in 12-well plates and co-transfected with 0.5 µg mini-Gs-NanoLuc and 0.5 µg KRAS- or Rab5-Venus, respectively, with either 0.5 µg SNAP-GLP-1R or SNAP-GIPR.

#### Mini-Gs-Venus recruitment NanoBRET assays

Mini-Gs-Venus was a gift from Prof Nevin Lambert, Augusta University, USA. INS-1 832/3 GLP-1R KO or GIPR KO cells were seeded in 12-well plates and co-transfected with 0.5 µg mini-Gs-Venus and either 0.5 µg GLP-1R- or GIPR-NanoLuc, respectively.

#### Nb37 bystander NanoBiT assays

The Nb37 assay constructs were kindly provided by Prof Asuka Inoue, Tohoku University, Japan. Nb37 (gene synthesized by GenScript with codon optimization) was C-terminally fused to SmBiT with a 15 amino acid flexible linker (GGSGGGGSGGSSSGGG), and the resulting construct referred to as Nb37-SmBiT. The C-terminal KRAS CAAX motif (SSSGGGKKKKKKSKTKCVIM) was N-terminally fused with LgBiT (LgBiT-CAAX). The Endofin FYVE domain (amino-acid region Gln739-Lys806) was C-terminally fused with LgBiT (Endofin-LgBiT). Gα_s_ (human, short isoform), Gβ1 (human), Gγ2 (human), and RIC8B (human, isoform 2) plasmids were inserted into pcDNA3.1 or pCAGGS expression plasmid vectors. INS-1 832/3 GLP-1R KO or GIPR KO cells were seeded in 6-well plates and co-transfected with 0.1 μg SNAP-GLP-1R or SNAP-GIPR, 0.5 μg Gα_s_, Gβ1, and Gγ2, 0.1 μg RIC8B, 0.1 μg CAAX-LgBiT or 0.5 μg Endofin-LgBiT with 0.1 μg or 0.5 μg Nb37-SmBiT, respectively, with 0.8 µg pcDNA3.1 added to the former to equalize DNA content.

All NanoBiT and NanoBRET readings were obtained in a Flexstation 3 plate reader. Briefly, 24 hrs after transfection, cells were detached, resuspended in NanoGlo Live Cell Reagent (Promega) with furimazine (1:20 dilution) and seeded into white 96-well half area plates. For NanoBiTs, baseline luminescence was recorded for 5 min at 37°C followed by 30 min with or without addition of GLP-1 or GIP at 100 nM for G protein and β-arrestin 2 recruitment assays, and at serial doses of up to 1 μM for the Nb37 bystander assays; readings were taken every 30 sec or every min, respectively. For NanoBRETs, baseline luminescent signals were recorded every min at 460 nm (NanoLuc emission peak) and 535 nm (Venus emission peak) over 5 min at 37 °C, followed by 30 min with or without the addition of 100 nM GLP-1 or GIP. Readings were normalized to well baseline and then to average vehicle-induced signal to establish the agonist-induced effect. AUCs from response curves were calculated for each agonist concentration and fitted to four-parameter curves using Prism 9 (GraphPad).

### Transfections

Transient transfection of plasmids was performed using Lipofectamine 2000 (Thermo Fisher) according to the manufacturer’s instructions. Experiments were performed 24 hrs after transfection unless otherwise indicated.

### Receptor degradation assays

#### High-content microscopy assay

The assay was adapted from a previous description [12]. INS-1 832/3 SNAP-GLP-1R or SNAP GIPR cells were seeded in complete medium in poly-D-lysine-coated black, clear-bottom plates. Once attached, cells were washed twice in PBS and incubated in fresh serum-free medium containing cycloheximide (50 µg/mL) to arrest protein translation. After 2 hrs, agonists were added in reverse time order (the longest time point being 8 hrs), with the medium replaced for the final 30 min of the experiment with complete medium containing 1 µM BG-OG to label total residual SNAP-GLP-1R or SNAP-GIPR. Wells were then washed 3X in HBSS and the microplate imaged by widefield microscopy, with quantification of total cellular receptor at each time point from segmented cell-containing regions as for the high-content internalization and recycling assays described above.

#### Degradation assays by immunoblotting

INS-1 832/3 SNAP-GLP-1R and SNAP-GIPR cells were seeded in 6-well plates (1.5 million per well) and cultured overnight prior to incubation in serum-free medium containing cycloheximide (50 µg/mL) for 2 hrs. Cells were then incubated with or without 100 nM GLP-1 or GIP for 6 hrs before being lysed in 1X TNE lysis buffer (20 mM Tris, 150 mM NaCl, 1 mM EDTA, 1% NP40, protease and phosphatase inhibitor cocktails) for 10 min at 4°C followed by cell scraping and sonication (3X, 10 secs each). The lysates were then frozen at -80°C for 2 min, thawed and centrifuged at 15,000 g for 10 min at 4°C. The supernatants were collected, fractionated by SDS-PAGE in urea loading buffer (200 mM Tris HCl pH 6.8, 5% w/v SDS, 8 M urea, 100 mM DTT, 0.02% w/v bromophenol blue) and analyzed by Western blotting. SNAP-GLP-1R and SNAP-GIPR were detected with an anti-SNAP-tag rabbit polyclonal antibody (P9310S, New England Biolabs, 1/1,000) followed by goat anti-rabbit HRP secondary (ab6271, Abcam, 1/2,000). Post-stripping, tubulin was labelled with anti-α-tubulin mouse monoclonal antibody (T5168, Sigma, 1/5,000) followed by sheep anti-mouse HRP secondary antibody (ab6721, Abcam, 1/5,000). Blots were developed with the Clarity Western enhanced chemiluminescence (ECL) substrate system (BioRad) in a Xograph Compact X5 processor and specific band densities quantified in Fiji.

### Measurement of receptor clustering by time-resolved fluorescence resonance energy transfer (TR- FRET)

The assay was performed as previously described [13]. INS-1 832/3 SNAP- GLP- 1R or SNAP-GIPR cells were labelled in suspension with 40 nM SNAP- Lumi4- Tb and 1 mM SNAP-Surface 649 (New England Biolabs, Hitchin, UK) for 1 hr at room temperature in complete medium. After washing, cells were resuspended in HBSS, and TR-FRET was monitored before and after addition of 100 nM GLP-1, GIP, or a mixture of GLP-1 and GIP at 37°C in a Spectramax i3x plate reader in HTRF mode. TR-FRET was quantified as the ratio of fluorescent signal at 665 nm to that at 616 nm, after subtraction of background signal at each wavelength.

### Raster image correlation spectroscopy (RICS)

INS-1 832/3 SNAP- GLP- 1R or SNAP-GIPR cells were seeded onto glass bottom MatTek dishes and surface-labelled with SNAP-Surface 488 (1 mM, 30 min at 37°C). After washing, cells were imaged at the basal plasma membrane in HBSS with 10 mM HEPES at 37°C either before or 5 min after stimulation with 100 nM GLP-1 or GIP, respectively. Time-lapse images of cells were acquired in a Zeiss LSM-780 inverted confocal microscope fitted with a 63x/1.2 NA water immersion objective. SNAP-Surface 488 was excited by a continuous wavelength laser at 488 nm and emission signal collected at 500–580 nm. The pinhole was set to one Airy unit. Optimized acquisition was performed to retrieve protein membrane diffusion values as described previously [14, 15]. Images of 256 × 256 pixels at 8-bit depth were collected using 80 nm pixel size and 5 μsec dwell time, for 250 consecutive frames. To characterize the waist of the point spread function (PSF), 200 frames of freely diffusing recombinant EGFP (20 mM) were continuously collected as described elsewhere [16, 17]. Analysis was performed on images where intensity traces were not decreased continuously by 20% or more over 50 frames to avoid possible bleaching artefacts that would interfere in diffusion coefficient measurements. A moving average (background subtraction) of 10 was applied, so that artefacts due to cellular motion or very slow-moving particles were avoided. The obtained 2D autocorrelation map was then fitted, and a surface map obtained with the characterized PSF and the appropriate acquisition values for line time and pixel time. Three different regions of interest (ROI) were analyzed within the same cell, with the corresponding regions drawn employing a 64 × 64-pixel square. RICS analysis was performed using the “SimFCS 4” software (Global Software, G-SOFT Inc., Champaign, Illinois) as described [18]. RICS analysis was performed in ROIs of 64 × 64 pixels at 4 random cytoplasmic areas per cell using a moving average (background subtraction) of 10 to discard possible artefacts due to cellular motion and slow-moving particles passing through. The autocorrelation 2D map was then fitted to obtain a surface map that was represented as a 3D projection with the residuals on top. As a rule, we focused on those regions with intensity fluctuation events in which the intensity changes were following short increasing or decreasing steps, avoiding abrupt intensity decays or increases.

### cAMP homogeneous time-resolved fluorescence (HTRF) assays

INS-1 832/3 cells were stimulated with increasing concentrations of GLP-1 or GIP, as well as with 10 μM forskolin to measure maximal responses, followed by lysis and cAMP HTRF immunoassay (cAMP Dynamic 2, 62AM4PEB, Cisbio, Codolet, France) according to the manufacturer’s instructions. Results were expressed as % forskolin responses and fitted to three-parameter curves using Prism 9 (GraphPad).

### Isolation and culture of pancreatic islets

For islet isolation, pancreata were infused via the common bile duct with RPMI-1640 medium containing 1 mg/mL collagenase from Clostridium histolyticum (Nordmark Biochemicals), dissected, and incubated in a water bath at 37°C for 10 min. Islets were subsequently washed and purified using a Histopaque gradient (Histopaque-1119, 11191, Sigma-Aldrich, and Histopaque-1083, 10831, Sigma-Aldrich). Isolated islets were allowed to recover overnight at 37°C in 5% CO_2_ in RPMI-1640 supplemented with 10% FBS and 1% penicillin/streptomycin.

### cAMP FRET assays

CAMPER reporter mice [19], with conditional expression of the cAMP FRET biosensor ^T^EPAC^VV^ [20], were purchased from Jackson Laboratory (Stock No: 032205) and crossed with Pdx1-Cre^ERT^ mice (in house) to generate mice with inducible ^T^EPAC^VV^ expression from pancreatic beta cells, used to isolate islets for *ex vivo* cAMP FRET assays. Isolated islets were treated overnight with 4-hydroxytamoxifen (4-OHT) to induce biosensor expression prior to Matrigel encasing on MatTek glass bottom dishes and imaging by FRET between CFP (donor) and YFP (acceptor) with CFP excitation and both CFP and YFP emission settings in a Zeiss LSM-780 inverted confocal laser-scanning microscope and a 20X objective to capture time-lapse recordings with image acquisition every 6 sec, and treatments manually added by pipetting. Specifically, islets were imaged in Krebs-Ringer Bicarbonate-HEPES (KRBH) buffer (140 mM NaCl, 3.6 mM KCl, 1.5 mM CaCl_2_, 0.5 mM MgSO_4_, 0.5 mM NaH_2_PO_4_, 2 mM NaHCO_3_, 10 mM HEPES, saturated with 95% O_2_/5% CO_2_; pH 7.4) containing 0.1% w/v BSA and 6 mM glucose (KRBH G6) for 1 min, then agonist at 100 nM was added and imaged for 10 min before addition of 10 μM forskolin + 100 μM isobutyl methylxanthine (IBMX) for the final 2 min of the acquisition to record maximal responses. Raw intensity traces for YFP and CFP fluorescence were extracted from whole islet ROIs using Fiji and YFP/CFP ratios calculated for each ROI and time-point. Responses were plotted relative to the average fluorescence intensity per islet during the 6 mM glucose baseline period, before agonist addition.

### Calcium assays

Imaging of INS-1 832/3 cells or whole-islet Ca^2+^ dynamics was performed as previously described [21]. Cells or Matrigel-encased islets from individual animals were loaded with the Ca^2+^ responsive dye Cal-520 AM (AAT Bioquest), pre-incubated for 1 hr in KRBH G6, and imaged in MatTek glass bottom dishes every 6 sec at 488 nm using a Nikon Eclipse Ti microscope with an ORCA-Flash 4.0 camera (Hamamatsu) and Metamorph software (Molecular Devices) while maintained at 37°C on a heated stage. Raw fluorescence intensity traces from cell-occupied areas or islet ROIs were extracted using Fiji. Responses were plotted relative to the average fluorescence intensity during the 6 mM glucose baseline period, before agonist addition.

### Insulin secretion assays

INS-1 832/3 cells were seeded in a 48-well plate and incubated in 3 mM glucose in full medium overnight before incubation with 11 mM glucose ± GLP-1/GIP at 100 nM in KRBH buffer containing 0.1% w/v BSA at 37°C. At the end of the treatments, the supernatant containing the secreted insulin was collected, centrifuged at 1,000 x g for 3 min, and transferred to a fresh tube. To determine total insulin content, cells were lysed using KRBH Buffer + 1% w/v BSA + 1% v/v Triton X-100 (Sigma). The lysates were sonicated 3 × 10 s in a water-bath sonicator and centrifuged at 10,000 x g for 10 min, and the supernatants collected. The samples were stored at -20°C until the insulin concentration was determined using an Insulin Ultra-Sensitive HTRF Assay kit (62IN2PEG, Cisbio, Codolet, France) according to the manufacturer’s instructions.

### Statistical analyses

All data analyses and graph generation were performed with GraphPad Prism 9.0. The statistical tests used are indicated in the corresponding figure legends. The number of replicates for comparisons represents biological replicates. Technical replicates within biological replicates were averaged prior to statistical tests. Data are represented as mean ± SEM. The p-value threshold for statistical significance was set at 0.05.

## Results

We first analyzed the trafficking characteristics of both receptors following stimulation with their cognate full length endogenous agonists, GLP-1(7-36)NH_2_ and GIP(1-42), using rat INS-1 832/3 beta cells in which the endogenous incretin receptor was deleted and the equivalent SNAP-tagged human receptor exogenously expressed (INS-1 832/3 SNAP-GLP-1R or SNAP-GIPR cells). Note that surface expression levels of SNAP-GLP-1R and SNAP-GIPR were similar within these two cell models (Supplementary Figure 1A). DERET assays, which detect disappearance of the receptor by a loss of TR-FRET signal between the receptor extracellular domain (ECD) and the extracellular buffer [13], revealed stark differences in the degree of internalization between the two receptors following stimulation with their native agonists, with the GLP-1R achieving around three times more internalization in the first hour post-stimulation with 100 nM GLP-1 compared to the GIPR for the same stimulation period with 100 nM GIP (Supplementary Figure 1B, C). Notably, there was negligible GLP-1R internalization in response to GIP, and vice versa, and no significant change to internalization for either receptor when using both agonists combined. Greater internalization of GLP-1R than of GIPR was observed across a wide concentration range (Figure 1A, Supplementary Figure 1D). Analysis of the rate of change of DERET signal indicated that GLP-1R endocytosis was significantly faster (Figure 1B). We corroborated these findings by high content microscopy analysis of receptor internalization in the same cells (Figure 1C), with significantly less internalization of GIPR compared to GLP-1R when stimulated with their respective endogenous agonists.

**Figure 1.**
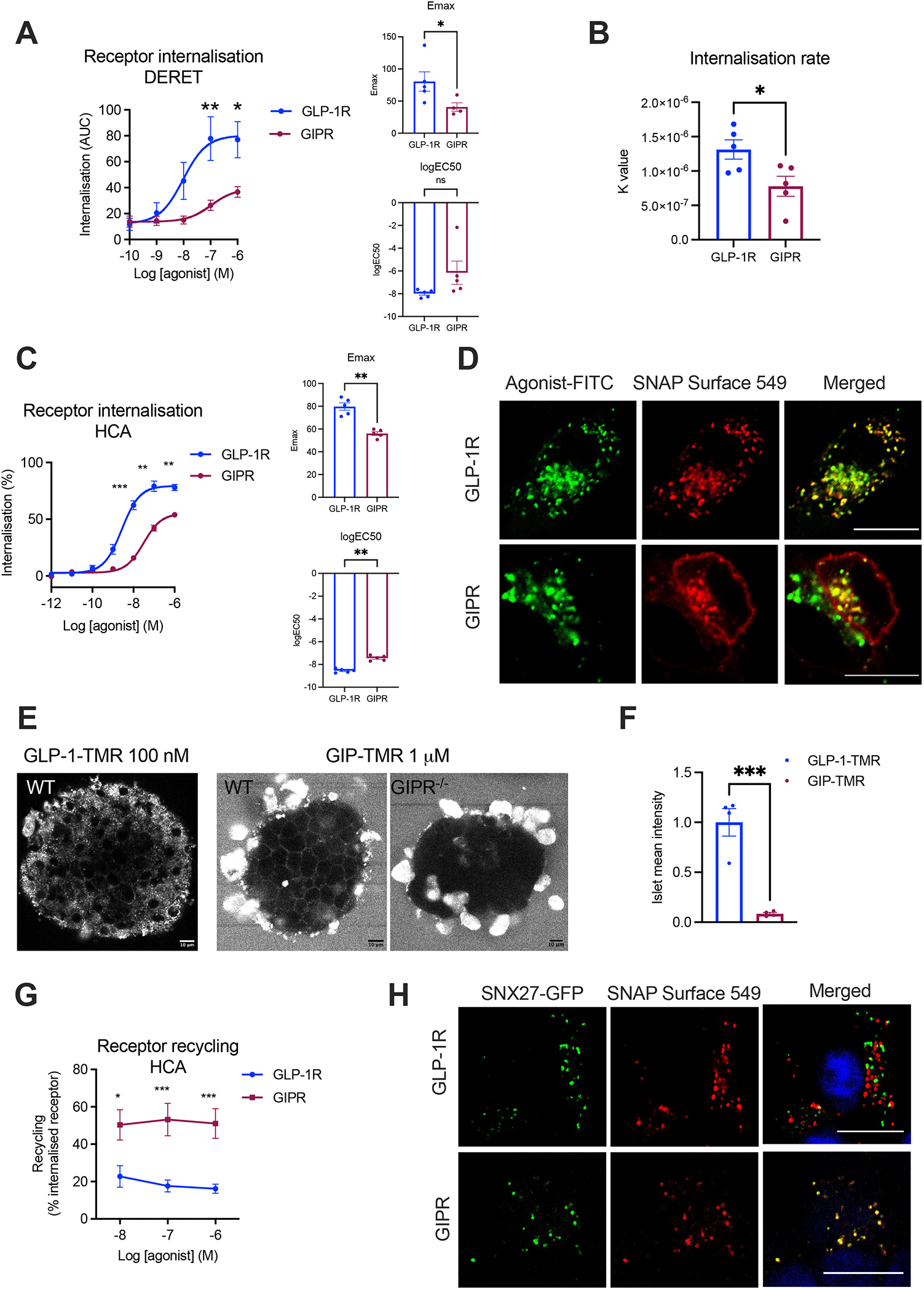
Beta cell GLP-1R *versus* GIPR trafficking patterns. (**A**) Internalization AUC dose response curves from DERET assays in INS-1 832/3 SNAP-GLP-1R *vs* SNAP-GIPR cells stimulated with the indicated concentrations of GLP-1 or GIP, respectively. Results were fitted to a 3-parameter dose response curve to obtain Emax and logEC50 for both receptors, with comparisons between these parameters included; *n*=5. (**B**) Rates of GLP-1R *vs* GIPR internalization (k values) derived from (A) by one-phase association (with Y0=0) of baseline-deleted DERET data; *n*=5. (**C**) GLP-1R *vs* GIPR internalization dose response curves measured by high-content microscopy assay (HCA) in the same cells as above. Results fitted and Emax and logEC50 comparisons included as above; *n*=5. (**D**) Confocal microscopy analysis of SNAP-GLP-1R *vs* SNAP-GIPR (red) localization following 30 min stimulation with 100 nM GLP-1-FITC or GIP-FITC, respectively, in the same cells as above. (**E**) Confocal microscopy analysis of isolated intact mouse islets stimulated with fluorescently labelled agonists as indicated: WT islets were imaged following stimulation with 100 nM GLP-1-TMR, while both WT and GIPR^-/-^ (KO) islets were imaged following stimulation with 1 µM GIP-TMR. (**F**) Quantification of surface GLP-1R *vs* GIPR levels in WT mouse islets using fluorescent agonist uptake data from (E); data corrected for binding affinity differences between both agonists; *n*=4. (**G**) GLP-1R *vs* GIPR recycling dose response curves measured by high-content microscopy assay (HCA) in cells from (C); *n*=5. (**H**) Confocal microscopy analysis of SNAP-GLP-1R *vs* SNAP-GIPR (red) co-localization with SNX27-GFP (green) following 3 hrs stimulation with 100 nM GLP-1 or GIP, respectively, in the same cells as above. Nuclei (DAPI), blue. Data are mean ± SEM, compared by paired or unpaired t-tests, or two-way ANOVA with Sidak’s post-hoc test; *p<0.05, **p<0.01, ***p<0.001; size bars: 10 µm.

In concordance with these results, 30 minutes of stimulation with the fluorescently labelled agonist GLP-1-FITC [22] resulted in the near complete co-internalization of SNAP-GLP-1R and fluorescent ligand, while the equivalent treatment with GIP-FITC led to only partial SNAP-GIPR endocytosis, with the receptor still clearly visible at the plasma membrane (Figure 1D, Supplementary Figure 1E). Moreover, when both receptors harboring different N-terminal tags (so that they could be differentially labelled) were expressed together in WT INS-1 832/3 cells, HALO-GLP-1R did again show faster internalization *versus* SNAP-GIPR following stimulation with a mixture of GLP-1 and GIP (Supplementary Figure 2). Finally, using alternative fluorescent conjugates labelled with TMR, the amount of TMR-labelled agonist uptake during the first 5 minutes of stimulation for each receptor correlated with the previously shown receptor internalization results in INS-1 832/3 SNAP-GLP-1R/SNAP-GIPR cell lines (Supplementary Figure 3A, B). Similar results were obtained in WT mouse primary islets, where GIP-TMR could be detected localized at the plasma membrane (albeit at low levels) while GLP-1-TMR was completely internalized after 15 minutes of stimulation (Figure 1E). Additionally, GIP-TMR signal at the plasma membrane was absent in islets from *Gipr*^*-/-*^ (KO) mice [23] labelled in parallel with the same concentration of GIP-TMR, demonstrating the specificity of labelling in WT islets. We quantified the reduction in TMR-labeled agonist binding for the GIPR *versus* the GLP-1R from data in Figure 1E (Figure 1F), which correlates with previously published RNAseq data indicating comparable higher levels of beta cell *Glp1rm versus Gipr* mRNA expression from mouse islets [24]. Additional experiments were performed using dispersed mouse islet cells, which also showed markedly reduced that GIP-TMR uptake compared to GLP-1-TMR (Supplementary Figure 3C).

We also analyzed the level of receptor recycling in INS-1 832/3 SNAP-GLP-1R or SNAP-GIPR cells by high content microscopy following stimulation with 100 nM GLP-1 or GIP, respectively, and detected significantly increased recycling rates for the GIPR compared to the GLP-1R (Figure 1G), an observation that correlated with sustained SNAP-GIPR, but not SNAP-GLP-1R, colocalization with the recycling factor SNX27 [25] fused to EGFP (Figure 1H). To determine the intracellular destination of internalized GLP-1Rs/GIPRs more precisely, we performed concentration response bystander NanoBRET assays using C-terminal NanoLuc-fused SNAP-tagged GLP-1R *versus* GIPR and either KRAS-Venus (plasma membrane) or Rab5-Venus (early endosome) co-expressed transiently in INS-1 832/3 cells (Figure 2). In these experiments, we again observed increased propensity for plasma membrane retention of the GIPR compared to the GLP-1R, with reduced maximal internalization responses but no changes in potency (Figure 2A). Agonist-mediated redistribution of GLP-1R to Rab5-positive early endosomes was clearly detectable, but virtually absent for the GIPR in this system (Figure 2B). In broad agreement with these results, using stable INS-1-SNAP-GLP-1R or -GIP cells, the majority of SNAP-GLP-1R signal could be detected in Rab5-Venus-positive endosomes after 10 minutes of stimulation with 100 nM GLP-1, while a sizeable amount of SNAP-GIPR still remained at the plasma membrane following stimulation with 100 nM GIP; internalized GIPRs nevertheless also localized to Rab5-Venus-positive endosomes (Supplementary Figure 4).

**Figure 2.**
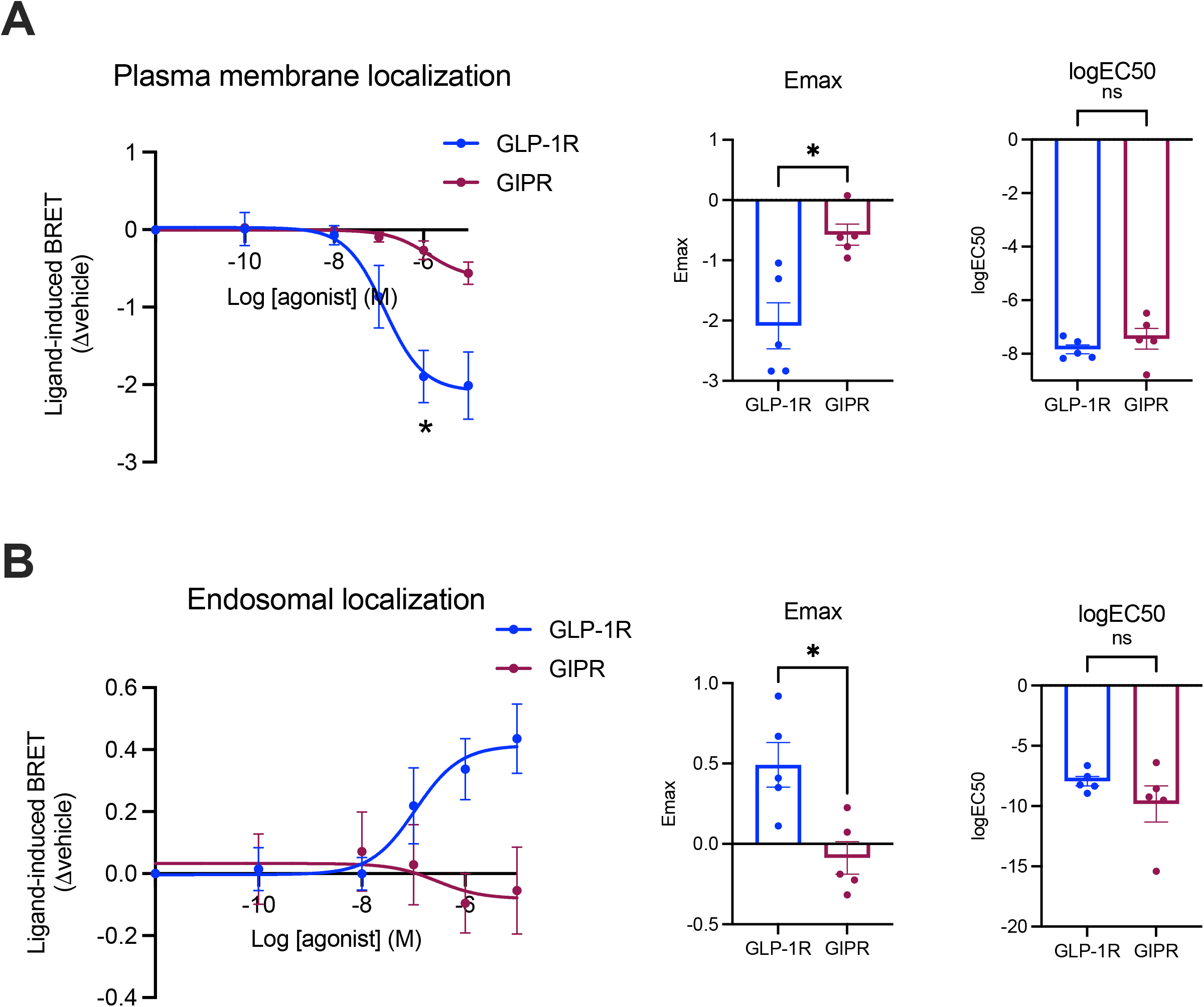
Endosomal *versus* plasma membrane localization of GLP-1R compared to GIPR in beta cells. (**A**) GLP-1R *vs* GIPR plasma membrane localization dose response curves from NanoBRET assays performed in INS-1 832/3 GLP-1R KO *vs* GIPR KO cells transiently expressing KRAS-Venus and GLP-1R- or GIPR-NanoLuc, after stimulation with the indicated concentrations of GLP-1 or GIP, respectively; results were fitted to 3-parameter dose response curves to obtain Emax and logEC50 for both receptors, with comparisons between these parameters included; *n*=5. (**B**) As for (A) but for GLP-1R *vs* GIPR endosomal localization dose response curves from NanoBRET assays performed in INS-1 832/3 GLP-1R KO *vs* GIPR KO cells transiently expressing Rab5-Venus and GLP-1R- or GIPR-NanoLuc, after stimulation with the indicated concentrations of GLP-1 or GIP, respectively; *n*=5. Data are mean ± SEM, compared by paired t-tests or two-way ANOVA with Sidak’s test; *p<0.05.

We have previously found that agonist-induced GLP-1R internalization is preceded by receptor clustering at the plasma membrane [13]. We therefore investigated whether the degree of clustering for each incretin receptor in a beta cell setting would reflect the differences observed in their internalization profiles (Figure 3). Employing raster imaging correlation spectroscopy (RICS) [16], we observed receptor clustering tendencies in INS-1 832/3 SNAP-GLP-1R or SNAP-GIPR cells labelled with the SNAP-Surface 488 probe under vehicle conditions as well as after 5 minutes of stimulation with 100 nM GLP-1 or GIP, respectively (Figure 3A). Quantification of receptor diffusion coefficients revealed that, while the GIPR exhibits slower basal diffusion, suggesting more clustering than the GLP-1R under vehicle conditions (a phenotype that correlates with our previously observed increased propensity for this receptor to segregate to cholesterol-rich lipid nanodomains under basal conditions [13]), agonist stimulation resulted in marked slowing of diffusion for both receptors (Figure 3B). Additionally, TR-FRET experiments suggested increased GLP-1R clustering when stimulated with GLP-1 (Figure 3C), an effect that could not be detected following GIP stimulation of the GIPR (Figure 3D). Moreover, clustering was not detectable when inverting the cognate agonists and no significant increases were detected by co-application of both agonists for each of the INS-1 832/3 receptor cell models (Figure 3C, D).

**Figure 3.**
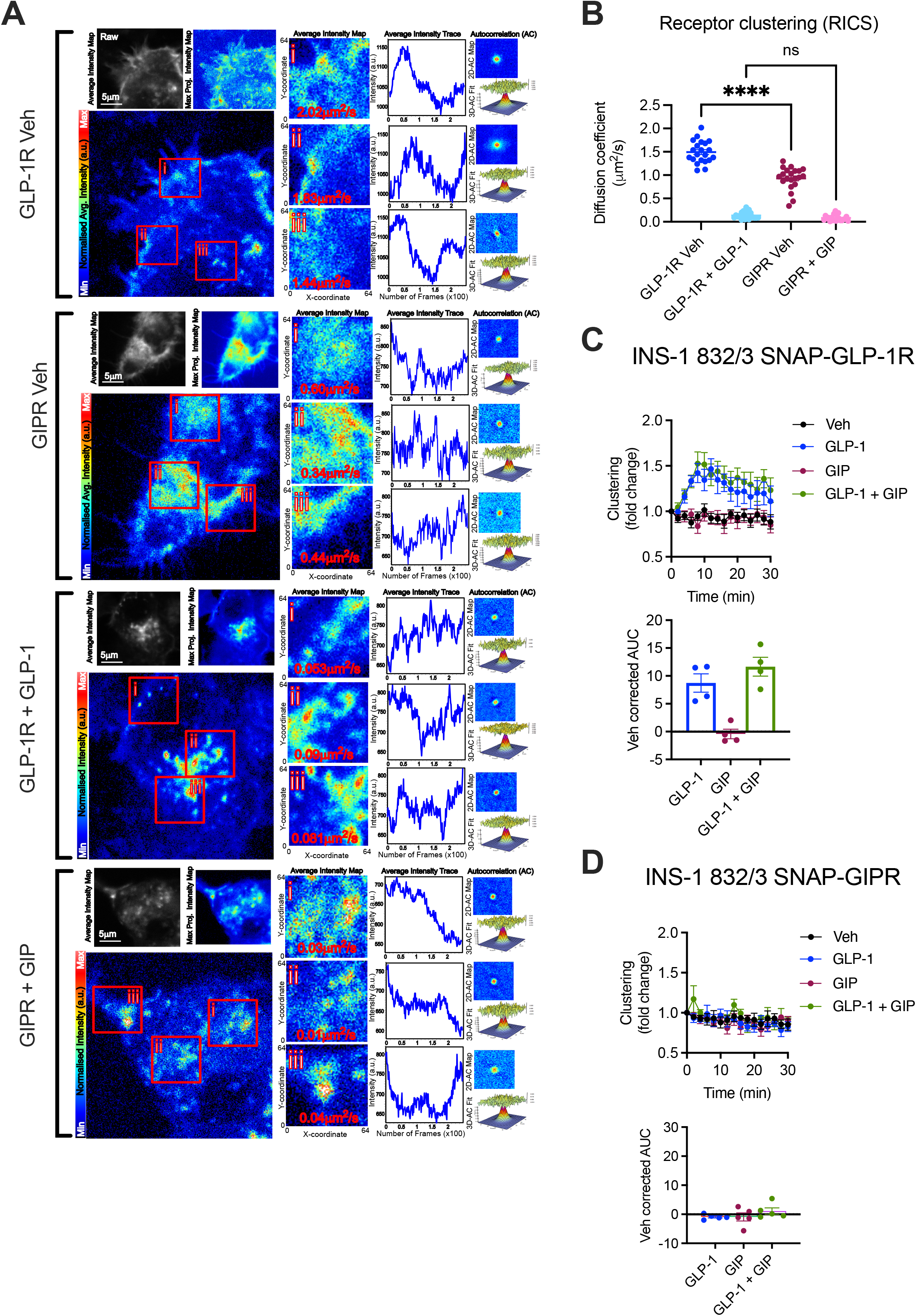
GLP-1R *versus* GIPR clustering propensities in beta cells. (**A**) Representative images from RICS analysis of GLP- 1R *vs* GIPR clustering, showing SNAP-Surface 488- labelled GLP-1Rs or GIPRs imaged in the basolateral plane of INS-1 832/3 SNAP-GLP-1R *vs* SNAP-GIPR cells after 5 min treatment in vehicle (Veh) and either 100 nM GLP-1 or GIP. Diffusion coefficients for individual ROIs are indicated on each image, with corresponding intensity traces, as well as 2D and fitted 3D autocorrelation maps for each ROI, are also depicted. (**B**) Average RICS diffusion coefficients for each receptor and treatment for each cell analyzed from *n*=4 experiments. (**C**) GLP- 1R clustering kinetics measured by TR-FRET in INS-1 832/3 SNAP-GLP-1R cells treated with 100 nM agonist as indicated, with vehicle-corrected AUCs included; *n*=4. (**D**) GIPR clustering kinetics measured by TR-FRET in INS-1 832/3 SNAP-GIPR cells treated with 100 nM agonist as indicated, with vehicle-corrected AUCs included; *n*=4. Data are mean ± SEM, compared by one-way ANOVA with Sidak’s test; ****p<0.0001; ns: non-significant.

We next analyzed the level of receptor degradation and lysosomal localization with a series of assays for both incretin receptors in INS-1 832/3 cells. We first assessed the total level of SNAP-GLP-1R *versus* SNAP-GIPR in INS-1 832/3 SNAP-GLP-1R or SNAP-GIPR cells in vehicle conditions or following 3-hour stimulation with 100 nM GLP-1 or GIP by Western blotting (Figure 4A, B). This experiment showed an increased propensity for degradation of the GLP-1R when compared to the GIPR. We next quantified the level of receptor degradation using a high content microscopy approach in which remaining total cellular SNAP-tag receptor is labelled after agonist incubation using the cell-permeable SNAP-tag probe BG-OG [12], and again observed faster receptor degradation for the GLP-1R *versus* the GIPR (Figure 4C). Finally, we also quantified the colocalization between each SNAP-tagged receptor and the lysosomes, finding significantly higher lysosomal targeting for the GLP-1R compared to the GIPR (Figure 4D, Supplementary Figure 5), a pattern that correlates with a reduced tendency for the GLP-1R *versus* the GIPR to localize to Rab11-positive recycling compartments following cognate agonist exposure (Supplementary Figure 6).

**Figure 4.**
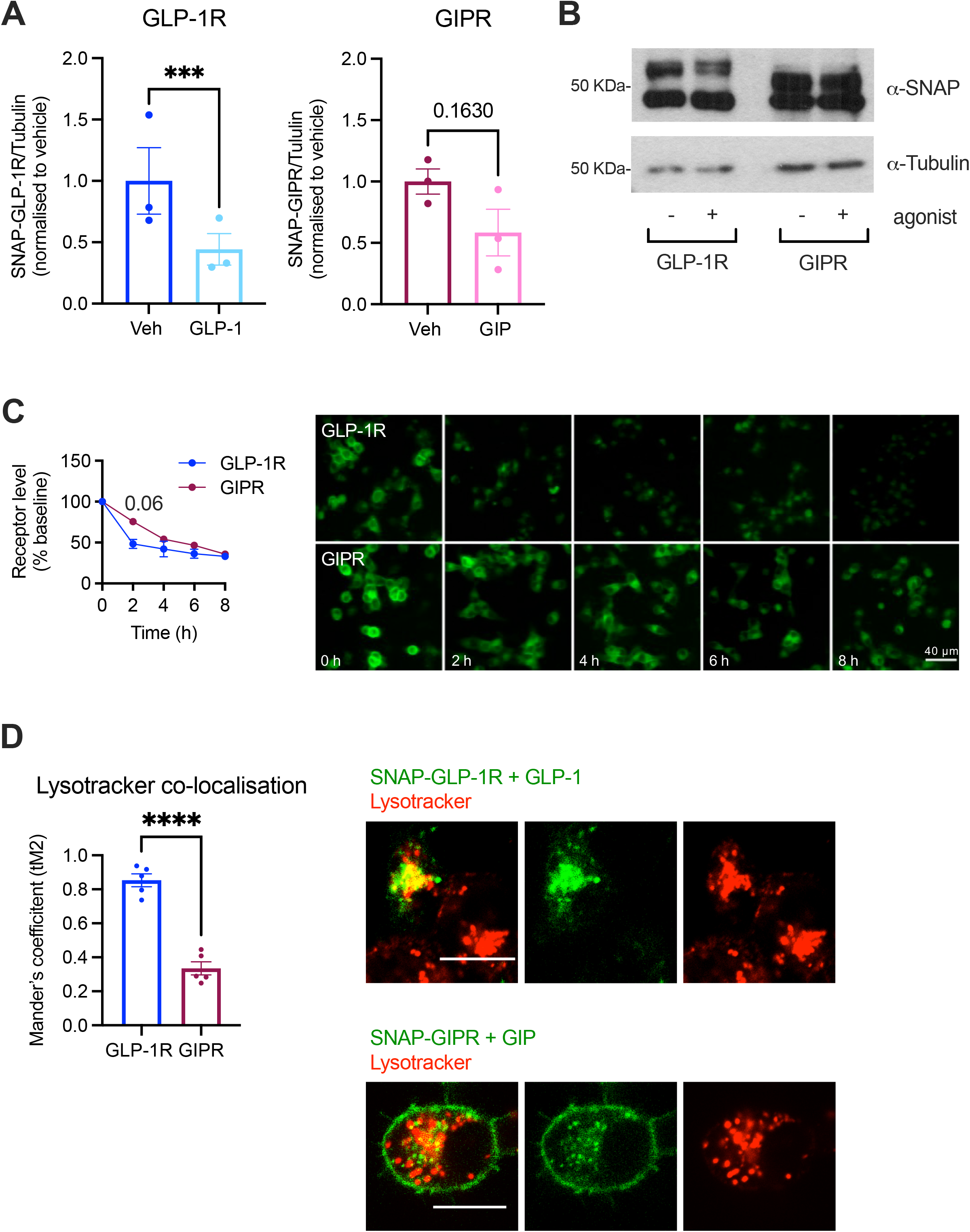
Beta cell GLP-1R *versus* GIPR degradation propensities. (**A**) Western blot assessment of SNAP-GLP-1R or GIPR over tubulin levels in INS-1 832/3 SNAP-GLP-1R or SNAP-GIPR cells with or without stimulation with 100 nM GLP-1 or GIP, respectively, for 6 hrs in the presence of the protein synthesis inhibitor cycloheximide; *n*=3. (**B**) Representative Western blot results from (A). Note that the top bands were used to quantify the SNAP-receptor levels, as they correspond to the glycosylated forms of the receptors, known to be biologically active and correctly inserted at the plasma membrane [13]. (**C**) Percentage of GLP-1R *vs* GIPR, labelled with the cell-permeable SNAP-tag probe BG-OG, and corresponding representative images from INS-1 832/3 SNAP-GLP-1R or SNAP-GIPR cells with or without stimulation with 100 nM GLP-1 or GIP, respectively, for the indicated times in the presence of cycloheximide; *n*=4. (**D**) Percentage of co-localization (Mander’s coefficient) and representative images of SNAP-GLP-1R *vs* -GIPR (labelled with SNAP-Surface 649) with Lysotracker Green in INS-1 832/3 SNAP-GLP-1R or SNAP-GIPR cells stimulated with 100 nM GLP-1 or GIP for 1 hr; *n*=5. Data are mean ± SEM, compared by ratio-paired or unpaired t-test or two-way ANOVA with Sidak’s post-hoc test; ***p<0.001, ****p<0.0001.

Having elucidated the main trafficking characteristics of both receptors, we next determined the coupling of each incretin receptor with specific signaling mediators, including Gα_s_, Gα_q_, Gα_i_ and β-arrestin 2 in INS-1 832/3 cells using NanoBiT complementation assays (Figure 5). As previously shown by our group using analogous assays in HEK293T cells [26], the GLP-1R was preferentially coupled to Gα_s_, followed by Gα_q_ and with minimum coupling to Gα_i_ proteins in response to GLP-1 stimulation (Figure 5A), while GIPR responses to GIP were markedly reduced for all readouts compared to GLP-1R (Figure 5B). As the values for Gα_s_ recruitment to the GIPR were almost as low as those obtained for Gα_i_ using this NanoBiT approach, we decided to verify whether the Gα_s_ results would be consistent when using a potentially more sensitive assay, namely a method based on NanoBRET between C-terminal NanoLuc-fused receptor and mini-Gs-Venus in the same cells as above (Figure 5C). Results with this method again showed significantly reduced Gα_s_ recruitment to the GIPR compared to the GLP-1R, although the difference between both receptors was less pronounced than previously found by NanoBiT complementation.

**Figure 5.**
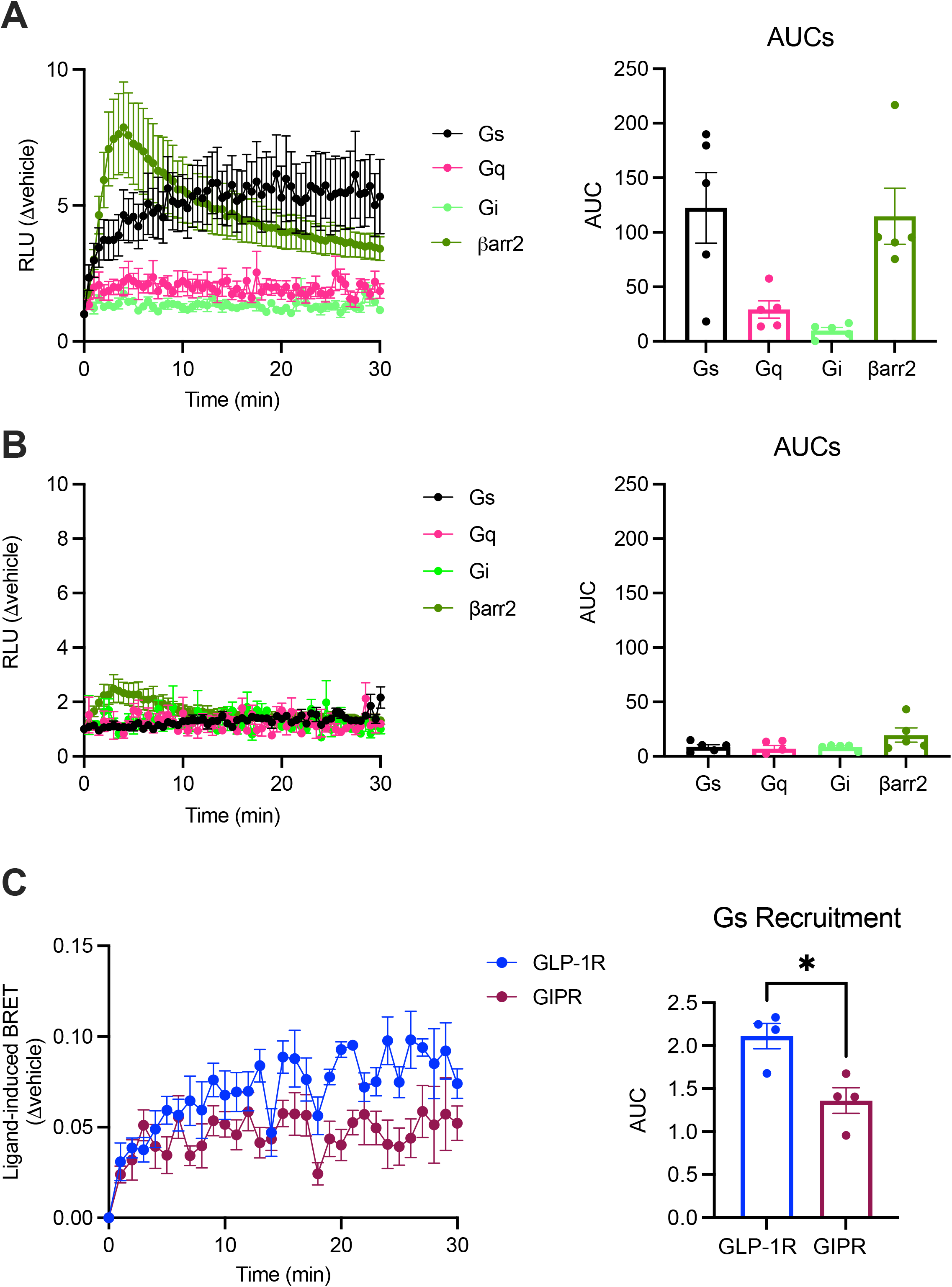
Beta cell GLP-1R *versus* GIPR G protein subtype and β-arrestin 2 recruitment characteristics. (**A**) Kinetics of Gα_s_, Gα_q_, Gα_i_, and β-arrestin 2 recruitment to the GLP-1R assessed by NanoBiT complementation assay in INS-1 832/3 GLP-1R KO cells transiently expressing GLP-1R-SmBiT and the corresponding mini-G protein subtype or β-arrestin 2. Responses to 100 nM GLP-1 normalized to vehicle and corresponding AUCs are shown; *n*=5. Kinetics of Gα_s_, Gα_q_, Gα_i_, and β-arrestin 2 recruitment to the GIPR assessed by NanoBiT complementation assay in INS-1 832/3 GIPR KO cells transiently expressing GIPR-SmBiT and the corresponding mini-G protein subtype or β-arrestin 2. Responses to 100 nM GIP normalized to vehicle and corresponding AUCs are shown; *n*=5. (**C**) NanoBRET assessment of GLP-1R *vs* GIPR recruitment of Gα_s_, performed in INS-1 832/3 GLP-1R KO *vs* GIPR KO cells transiently expressing mini-Gs-Venus and either GLP-1R- or GIPR-NanoLuc, after stimulation with 100 nM GLP-1 or GIP, respectively, with corresponding AUCs also shown; *n*=4. Data are mean ± SEM, compared by paired t-test; ***p<0.05.

Next, to determine the spatiotemporal pattern of Gα_s_ activation elicited by the two receptors, we employed a recently developed bystander NanoBiT signaling assay based on the recruitment of activated Gα_s_-recognizing nanobody 37 (Nb37) to plasma membrane and endosomal locations in response to specific agonist stimulations [27] (Figure 6, Supplementary Figure 7A, B). While the assay showed no significant differences in the recruitment of Nb37 to active GLP-1Rs or GIPRs within the plasma membrane (Figure 6A, Supplementary Fig 7A), there was a clear difference in GLP-1R *versus* GIPR endosomal activity, including a profoundly reduced GIPR Emax response despite increased potency for endosomal signaling with this receptor compared to the GLP-1R (Figure 6B, Supplementary Figure 7B). We also employed a different approach to quantify plasma membrane and endosomal activity from both incretin receptors based on a NanoBRET assay to measure the recruitment of NanoLuc-fused mini-Gs proteins to either KRAS- or Rab5-Venus constructs (Supplementary Figure 7C). We have recently shown that co-expression of mini-Gs with a cognate GPCR results in the inhibition of the normal level of receptor internalization in response to agonist stimulation [28]. For this reason, we were only able to perform this assay by using supra-physiological levels (100 nM) of GLP-1 and GIP and we could not obtain concentration response curves. We were nevertheless able to show a more persistent level of plasma membrane mini-Gs recruitment for GIPR when compared to the GLP-1R, which correlates with the above shown internalization patterns for each receptor. Endosomal mini-Gs recruitment, indicating the presence of active receptors in this intracellular compartment, could only be detected following a 30-minute pre-stimulation period with GLP-1 or GIP, again most likely due to the delayed endocytosis caused by the co-expression of mini-Gs protein. Nevertheless, active receptors were detectable for the 30-to 60-minute stimulation period and, again in agreement with our trafficking results, levels were significantly reduced for the GIPR when compared to the GLP-1R.

**Figure 6.**
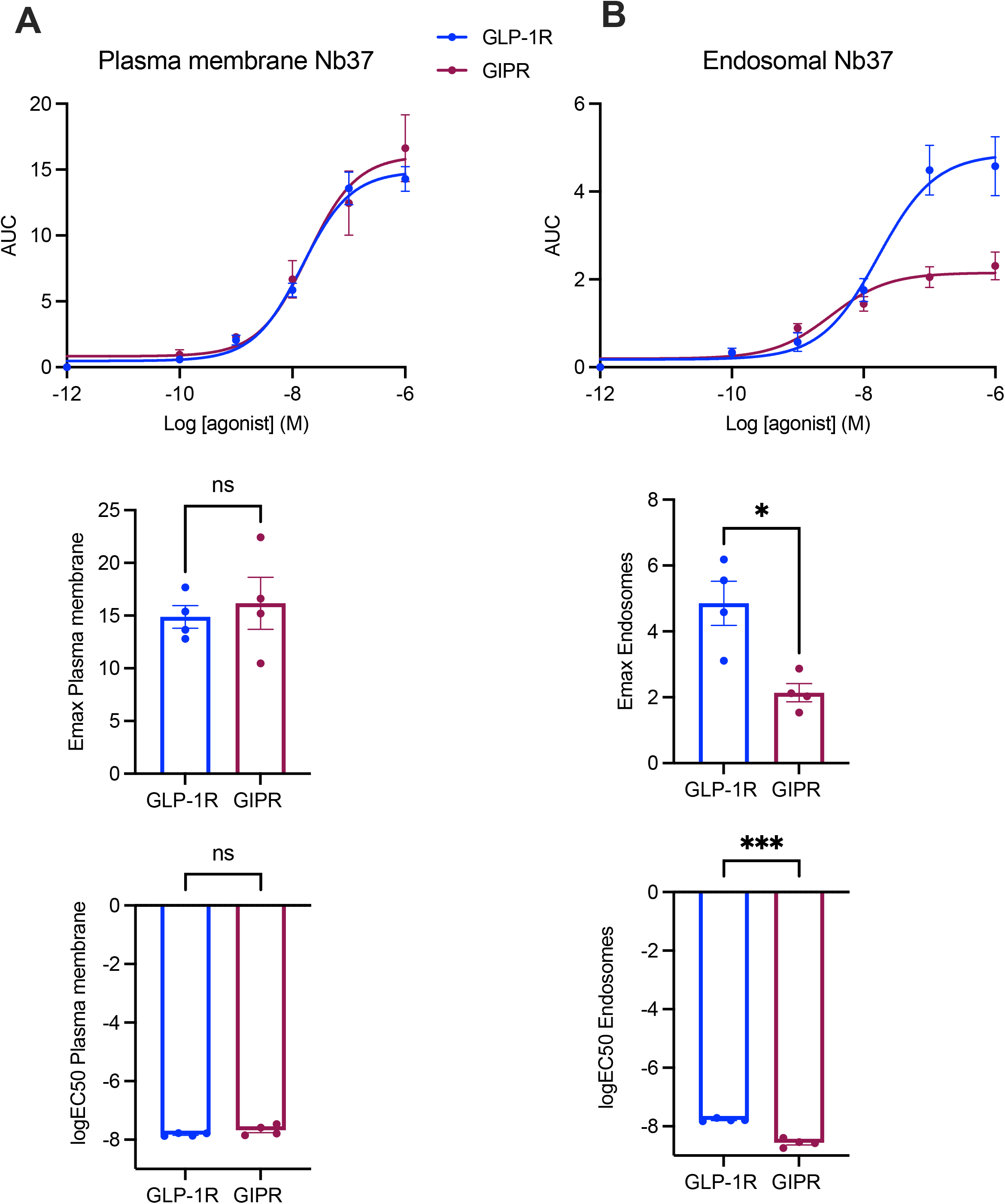
GLP-1R *versus* GIPR endosomal *versus* plasma membrane activity in beta cells. (**A**) GLP-1R *vs* GIPR plasma membrane activity dose response curves from bystander NanoBiT signaling assays performed in INS-1 832/3 GLP-1R KO *vs* GIPR KO cells transiently expressing Nb37-SmBiT, LgBiT-CAAX and SNAP-GLP-1R or -GIPR, after stimulation with the indicated concentrations of GLP-1 or GIP, respectively; results were fitted to 3-parameter dose response curves to obtain plasma membrane Emax and logEC50 for both receptors, with comparisons between these parameters included; *n*=4. (**B**) As for (A) but for GLP-1R *vs* GIPR endosomal activity in INS-1 832/3 GLP-1R KO *vs* GIPR KO cells transiently expressing Nb37-SmBiT, Endofin-LgBiT and SNAP-GLP-1R or -GIPR, after stimulation with the indicated concentrations of GLP-1 or GIP, respectively; *n*=4. Data are mean ± SEM, compared by paired t-tests; *p<0.05, ***p<0.001.

Finally, we determined the downstream signaling effects of both incretin receptors when stimulated with their native cognate agonists in beta cells (Figure 7), and found a significant shift towards a more potent cAMP response for the GIPR *versus* the GLP-1R in INS-1 832/3 cells (Figure 7A), which, for the case of primary mouse islet beta cells, resulted in a non-significant increase in cAMP response to 100 nM GIP compared to GLP-1 stimulations (Figure 7B). For intracellular calcium influx in INS-1 832/3 cells, while there was an initial trend for a decreased GIPR compared to GLP-1R response during the first minute of stimulation, this tendency was reversed for the following 4 minutes of agonist exposure, resulting in a zero net difference between both receptors (Figure 7C). However, in primary islets, there was a significant increase in calcium influx in response to GIP *versus* GLP-1 stimulations (Figure 7D). Finally, insulin secretion assays performed in INS-1 832/3 cells showed a non-significant tendency towards improvement following GIP compared to GLP-1 exposure, with responses being specific for either the GIPR (for GIP) or the GLP-1R (for GLP-1), as they were abolished in the absence of each receptor in the corresponding GLP-1R or GIPR KO cells (Figure 7E).

**Figure 7.**
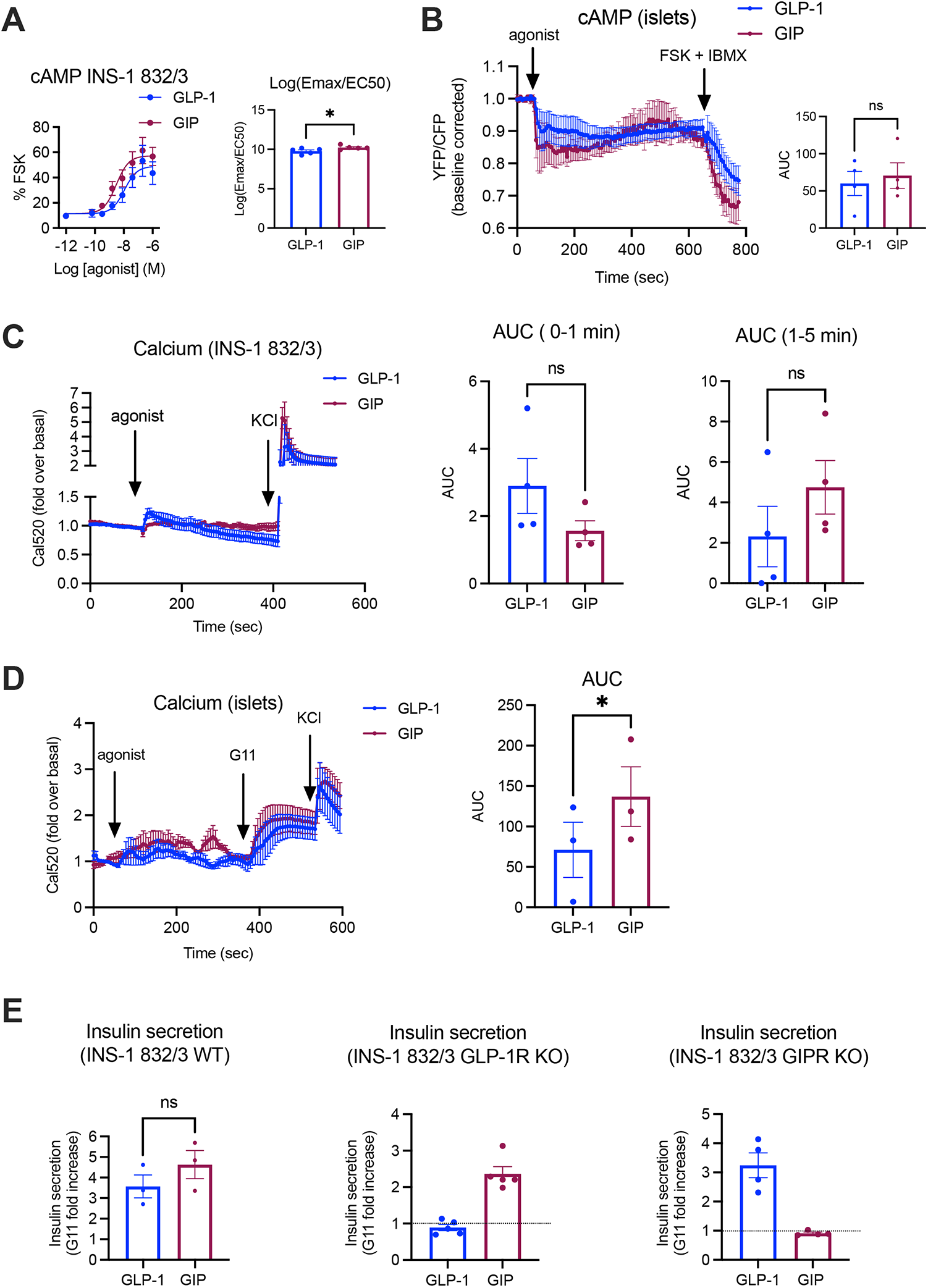
Functional analysis of GLP-1R *versus* GIPR signaling in beta cells. (**A**) cAMP dose response curves to GLP-1 *vs* GIP assessed in INS-1 832/3 cells by HTRF assay; results were fitted to 3-parameter dose response curves to obtain plasma membrane Emax and logEC50 for both receptors, depicted here combined as log(Emax/EC50) for each receptor; *n*=5. (**B**) cAMP FRET responses to 100 nM GLP-1 or GIP from isolated and 4-OHT-treated Pdx1-Cre^ERT^/CAMPER mouse islets; including agonist AUCs calculated for each receptor; *n*=4. (**C**) INS-1 832/3 calcium responses (using the calcium indicator Cal520-AM) to 100 nM GLP-1 or GIP, including agonist AUCs calculated for 0-1 min and 1-5 min responses for each receptor; *n*=4. (**D**) Calcium responses to 100 nM GLP-1 *vs* GIP from purified WT mouse islets loaded with Cal520-AM, including agonist AUCs for each receptor; *n*=4. (**E**) Insulin secretion (fold increase to 11 mM glucose) responses to 100 nM GLP-1 *vs* GIP from INS-1 832/3 WT (*n*=3), GLP-1R KO (*n*=5) and GIPR KO (*n*=4) cells. Data are mean ± SEM, compared by paired t-tests; *p<0.05, ns: non-significant.

## Discussion

In this study, we have established the main pattern of spatiotemporal signaling for each incretin receptor in a relevant cellular system, primarily INS-1 832/3 rat beta cells (with selected assays in primary mouse islets), where the receptors are expressed endogenously. We have found marked differences in the trafficking and signaling characteristics from the two receptors, with GIPR associated with significantly reduced internalization and degradation propensities but increased recycling when compared to GLP-1R by several different techniques, a pattern that correlates with reduced activity from endosomes but no significant differences in plasma membrane receptor activity. While the trafficking of the GIPR has been examined before in heterologous HEK293/HEK293T cells, this is to our knowledge the first in-depth examination of these patterns in pancreatic beta cells. Interestingly, while the net effect of these trafficking variations appears to be an increased level of sustained GIPR localization at the plasma membrane, we were still able to detect fluorescent GIP-FITC intracellularly after 30 minutes of agonist stimulation (although uptake of GIP-TMR, as opposed to that of GLP-1-TMR, was negligible at 5 minutes post-stimulation, suggesting reduced agonist internalization rates for the GIPR *versus* the GLP-1R). This observation points towards active GIPRs shuttling in and out of cells by a slow internalizing but rapidly recycling pathway, depositing their agonist in an intracellular location prior to receptor recycling, in a mechanism reminiscent to that previously observed by us for the related glucagon receptor (GCGR) [27].

A previous study from our group performed in HEK293T cells also reported reduced internalization propensity for the GIPR *versus* the GLP-1R in response to stimulation with their corresponding native agonists, a pattern that correlated with reduced recruitment of β-arrestin 2 to the GIPR [26], an effect already observed before in a prior study from a separate group performed in HEK293 cells [29]. Here, we again observe reduced propensity for β-arrestin 2 recruitment by the GIPR, this time in a beta cell context. The role of β-arrestins on incretin receptor trafficking and signaling has previously been investigated by us from several angles, for example by the use of biased agonists with different capabilities for β-arrestin recruitment [8, 26], using *in vivo* conditional β-arrestin 2 knockout mouse models [21], or with *in vitro* cell systems with deleted β-arrestin 1/2 expression [13, 26]. In all these instances, β-arrestin recruitment closely correlated with the degree of incretin receptor internalization, but alterations in β-arrestin expression levels or complete β-arrestin deletion did not lead to significant effects in receptor endocytosis, but rather resulted in the prolongation of cAMP/PKA signaling duration, suggesting that the main effect of this important signaling mediator lies in the steric hindrance caused by its binding to the receptor, leading to reduced access of Gα_S_ to its binding pocket and promoting homologous receptor desensitization [30]. Of note, similar reduced GIPR *versus* GLP-1R β-arrestin recruitment and internalization into a Rab5-positive endosomal compartment were also apparent in a separate study [31], although in this instance the authors focused on the comparison between responses from single and dual agonists such as tirzepatide or MAR709 for each receptor rather than performing a direct comparison of both receptor responses. Also interestingly, we previously observed that *in vivo* GIPR responses were less affected by β-arrestin 2 deletion specifically from pancreatic beta cells [21], suggesting a reduced reliance on β-arrestin 2 to regulate GIPR signaling effects in the pancreas.

Despite the abovementioned effects, the GIPR seems not only to be associated with reduced β-arrestin 2 recruitment, but paradoxically with a general dampening in the recruitment of other downstream effectors including Gα_s_ and Gα_q_ proteins, matching our previous observations in HEK293T cells [26]. While the reasons behind this effect are not fully elucidated, it is important to point out that the GIPR has previously been found to have significantly higher levels of basal activity *versus* the GLP-1R [29]. Accordingly, we have previously found increased basal association of this receptor with cholesterol-rich plasma membrane nanodomains [13], which are signaling hotspots rich in G proteins [32]. Consistent with this observation, we found in the present study significantly reduced GIPR *versus* GLP-1R basal diffusion rates, indicative of increased basal clustering, which correlated with an overall decrease in clustering fold increases following GIPR stimulation, again suggesting increased basal activity for the GIPR when compared to the GLP-1R. This observation might potentially contribute to explain the disconnect between the overall Gα_s_ and Gα_q_ levels of recruitment to the GIPR and the observed tendency towards increased cAMP, calcium and insulin secretion responses to GIP *versus* GLP-1 in INS-1 832/3 cells. Also of note, these tendencies are present despite the measured loss of recruitment of active Gα_s_ to endosomal compartments, suggesting that, as previously observed by us for the GLP-1R when stimulated with biased compounds affecting its capacity for endosomal localization and activity [8, 33], the plasma membrane is the main contributor to the overall signaling output of incretin receptors. This also suggests the existence of increased mechanisms of signal amplification associated with the GIPR, potentially related to differential interactions with downstream signaling mediators *versus* the GLP-1R. It is worth noting, however, that direct comparisons between different recruitment and activity assays are problematic due to potential differences in receptor *versus* effector expression levels leading to variations in stoichiometry as well as different dynamic ranges for each specific assay which might complicate their interpretation. Nevertheless, it is interesting to note that the GIPR also appears to signal more prominently than the GLP-1R in primary islets, where we measured a non-significant tendency towards increased cAMP and a significant increase in calcium responses to GIP *versus* GLP-1 despite a 12-fold reduction in endogenous surface GIPR *versus* GLP-1R levels estimated in WT mouse islets in this study by quantification of TMR-labelled agonist uptake.

In summary, we have described here significant differences in the trafficking and spatiotemporal signaling propensities of the two incretin receptors following their stimulation with native agonists in beta cells. While the GLP-1R is a rapidly internalizing receptor with increased propensity for β-arrestin 2 recruitment, endosomal localization and activity, and lysosomal degradation, the GIPR is associated with reduced coupling to G proteins and β-arrestin 2, as well as reduced internalization and endosomal activity, increased recycling and an overall increase in beta cell signaling despite highly reduced levels of endogenous surface receptor expression. These characteristics suggest that GLP-1R and GIPR signaling from beta cells are differentially regulated, highlighting the rationale for the development of dual agonists eliciting complementary beneficial effects from each receptor, as exemplified by the successful clinical development of the dual GLP-1R – GIPR agonist tirzepatide [6], a ligand which combines reduced internalization and β-arrestin 2 recruitment at the GLP-1R with GIPR stimulation [31].

## Supplementary Figure Legends

**Supplementary Figure 1. Beta cell GLP-1R *versus* GIPR trafficking – extra data**. (**A**) SNAP-GLP-1R *vs* SNAP-GIPR cell surface levels in in INS-1 832/3 SNAP-GLP-1R *vs* SNAP-GIPR cells. (**B**) Level of internalization of GLP-1R compared to GIPR, measured by DERET assay in INS-1 832/3 SNAP-GLP-1R *vs* SNAP-GIPR cells following 1 hr stimulation with 100 nM GLP-1, GIP or a combination of GLP-1 and GIP and expressed as response AUC; *n*=3. TR-FRET traces from (B). (**D**) TR-FRET traces from Figure 1A. (**E**) Amino acid sequences indicating the position of the modifications in the FITC-labelled agonists.

**Supplementary Figure 2. SNAP-GIPR/HALO-GLP-1R internalization in beta cells**. Time frame montage with the indicated time points of an INS-1 832/3 WT cell co-expressing SNAP-GIPR + HALO-GLP-1R prior to labeling with SNAP-Surface 549 + HaloTag AlexaFluor 660 impermeable probes and imaging at 10 sec/frame for 10 min post-stimulation with a mixture of 100 nM GLP-1 + GIP; size bar: 10 µm.

**Supplementary Figure 3. GLP-1-*versus* GIP-TMR uptake in beta cells**. (**A**) Amino acid sequences indicating the position of the modifications in the TMR-labelled agonists. (**B**) Time frame montage with the indicated time points from INS-1 832/3 SNAP-GLP-1R (top) *vs* -GIPR (bottom) cells labeled with SNAP-Surface 649 and stimulated with 100 nM GLP-1- or GIP-TMR for 5 min; size bars: 10 µm. (**C**) Quantification of TMR uptake in INS-1 832/3 WT cells treated with vehicle or 100 nM GLP-1-*vs* GIP-TMR +/-blocking with the corresponding unlabeled agonist at 10 µM.

**Supplementary Figure 4. GLP-1R *versus* GIPR localization to early endosomes in beta cells**. (**A**) Time frame montage with the indicated time points from INS-1 832/3 SNAP-GLP-1R cells transfected with Rab5-Venus prior to labeling with SNAP-Surface 549 and stimulation with 100 nM GLP-1 for 5 min. (**B**) As in (A) but from INS-1 832/3 SNAP-GIPR cells transfected with Rab5-Venus prior to labeling with SNAP-Surface 549 and stimulation with 100 nM GIP for 5 min. Size bars: 10 µm.

**Supplementary Figure 5. GLP-1R *versus* GIPR localization to lysosomes in beta cells**.

(**A**) Time frame montage with the indicated time points from INS-1 832/3 SNAP-GLP-1R cells labeled with Lysotracker Green + SNAP-Surface 649 and stimulation with 100 nM GLP-1 for 10 min. (**B**) As in (A) but from INS-1 832/3 SNAP-GIPR cells labeled with Lysotracker Green + SNAP-Surface 649 and stimulation with 100 nM GIP for 15 min. Size bars: 10 µm.

**Supplementary Figure 6. GLP-1R *versus* GIPR localization to recycling endosomes in beta cells**. (**A**) Time frame montage with the indicated time points from INS-1 832/3 SNAP-GLP-1R cells transfected with Rab11-Venus prior to labeling with SNAP-Surface 549 and stimulation with 100 nM GLP-1 for 5 min. (**B**) As in (A) but from INS-1 832/3 SNAP-GIPR cells transfected with Rab11-Venus prior to labeling with SNAP-Surface 549 and stimulation with 100 nM GIP for 5 min. Size bars: 10 µm.

**Supplementary Figure 7. GLP-1R *versus* GIPR endosomal *versus* plasma membrane activity in beta cells – extra data**. (**A**) Bystander NanoBiT plasma membrane response curves for each agonist concentration from dose response curves shown in Figure 6A. (**B**) Bystander NanoBiT endosomal response curves for each agonist concentration from dose response curves shown in Figure 6B. (**C**) Recruitment of mini-Gs-NanoLuc to plasma membrane (KRAS-Venus) or endosomal (Rab5-Venus) locations, measured by NanoBRET assay in INS-1 832/3 GLP-1R KO *vs* GIPR KO cells transiently transfected with SNAP-GLP-1R or -GIPR, and stimulated with 100 nM GLP-1 or GIP, respectively; corresponding AUCs are also shown; *n*=4 for plasma membrane and *n*=6 for endosomal responses (measured following a 30 minute pre-stimulation period). Data are mean ± SEM, compared by paired t-tests; *p<0.05, **p<0.01.

## Acknowledgements

This work was supported by MRC grant number MR/R010676/1 to A.T. and B.J., and by UKRI COVID-19 Grant Extension Allocation (coA) to A.T. A.T. also acknowledges further support from the EFSD, Diabetes UK, Eli Lilly, the Commonwealth, and the Integrated Biological Imaging Network (IBIN). The authors thank Dr Aida Martinez-Sanchez, Imperial College London, UK, for providing animal project license access and Dr Kyle Sloop, Eli Lilly and Company, Indianapolis, Indiana, USA, for the *Gipr*^*-/-*^ mice.

