## Supplementary Figure 1 for "An examination of the divergent spatiotemporal signaling of GLP-1R *versus* GIPR in pancreatic beta cells"

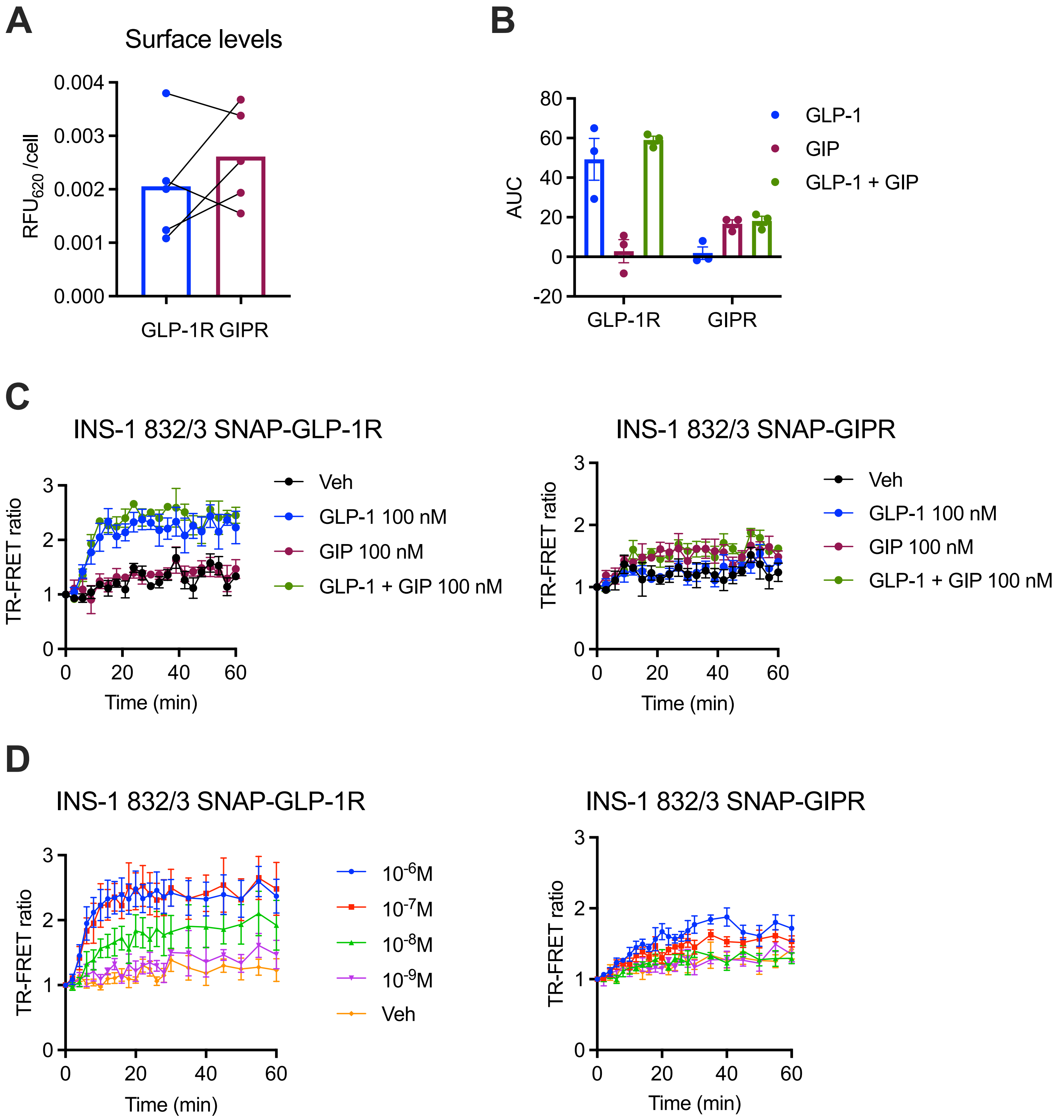


## E

| Shorter name | Peptide | 1 | 2 | 3 | 4 | 5 | 6 | 7 | 8 | 9 | 10 | 11 | 12 | 13 | 14 | 15 | 16 | 17 | 18 | 19 | 20 | 21 | 22 | 23 | 24 | 25 | 26 | 27 | 28 | 29 | 30 | 31 | 32 | 33 | 34 | 35 | 36 | 37 | 38 | 39 | 40 | 41 | 42 |
|---|---|---|---|---|---|---|---|---|---|---|---|---|---|---|---|---|---|---|---|---|---|---|---|---|---|---|---|---|---|---|---|---|---|---|---|---|---|---|---|---|---|---|---|
| GLP-1-FITC | GLP-1(7-36)-K28-FITC) | His | Ala | Glu | Gly | Thr | Phe | Thr | Ser | Asp | Val | Ser | Ser | Tyr | Leu | Glu | Gly | Gln | Ala | Ala | Lys | Glu | Phe | Ile | Ala | Trp | Leu | Val | Lys-FITC | Gly | Arg | NH2 | | | | | | | | | | | |
| GIP-FITC | GIP(1-30)-K30-FITC | Tyr | Ala | Glu | Gly | Thr | Phe | Ile | Ser | Asp | Tyr | Ser | Ile | Ala | Met | Asp | Lys | Ile | His | Gln | Gln | Asp | Phe | Val | Asn | Trp | Leu | Leu | Ala | Gln | Lys-FITC | | | | | | | | | | | | |
