## Supplementary figures and images for "An examination of the divergent spatiotemporal signaling of GLP-1R *versus* GIPR in pancreatic beta cells"

### Supplementary Figure 2

#### Supplementary Figure 2

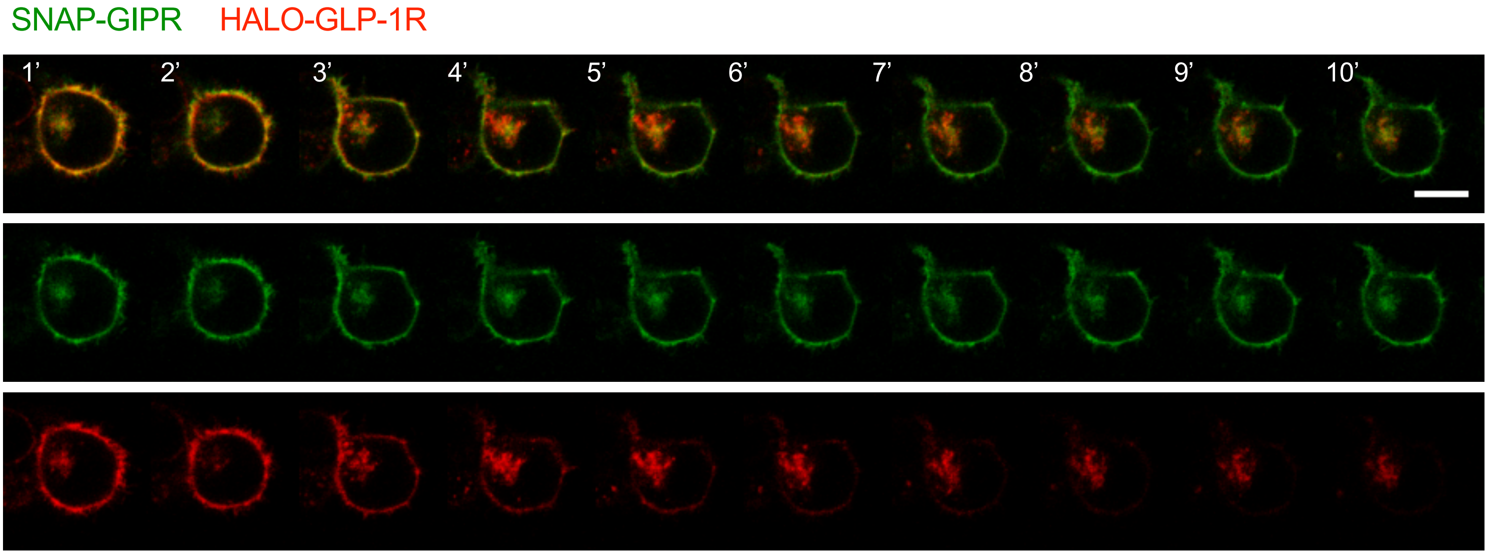

### Supplementary Figure 4

#### Supplementary Figure 4

## A

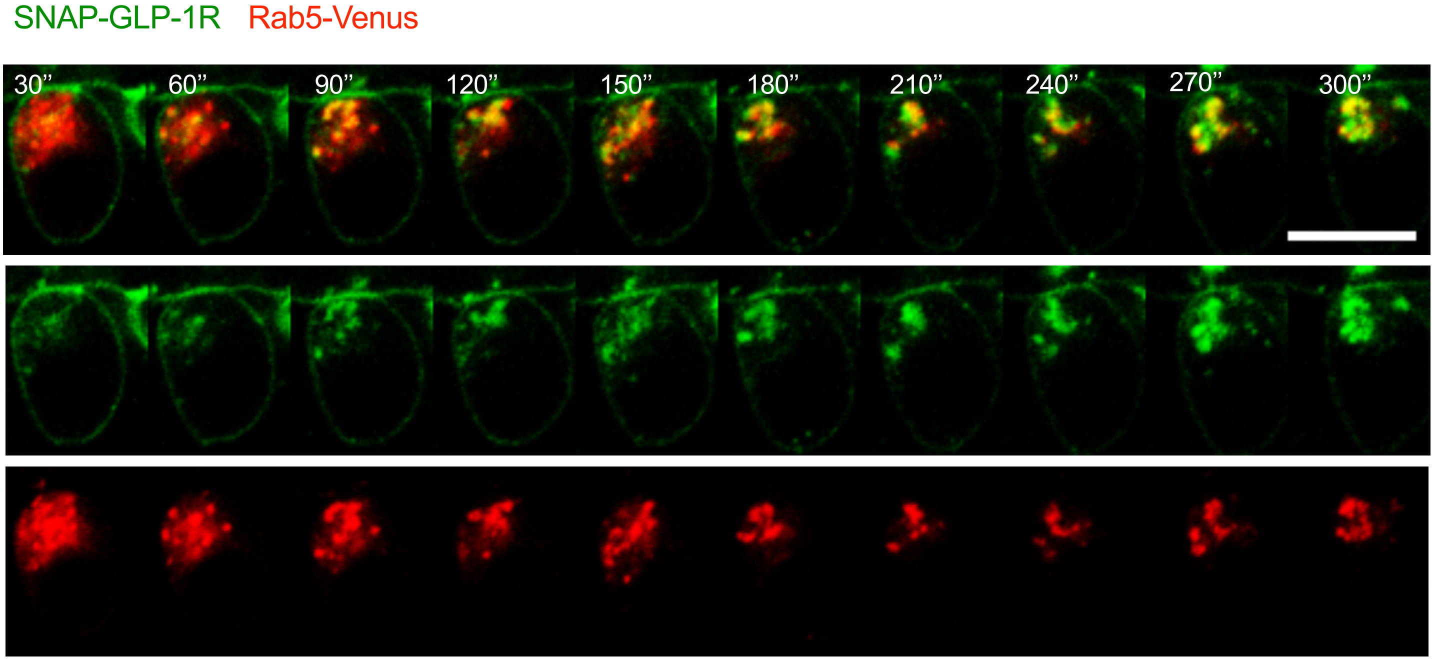


## B

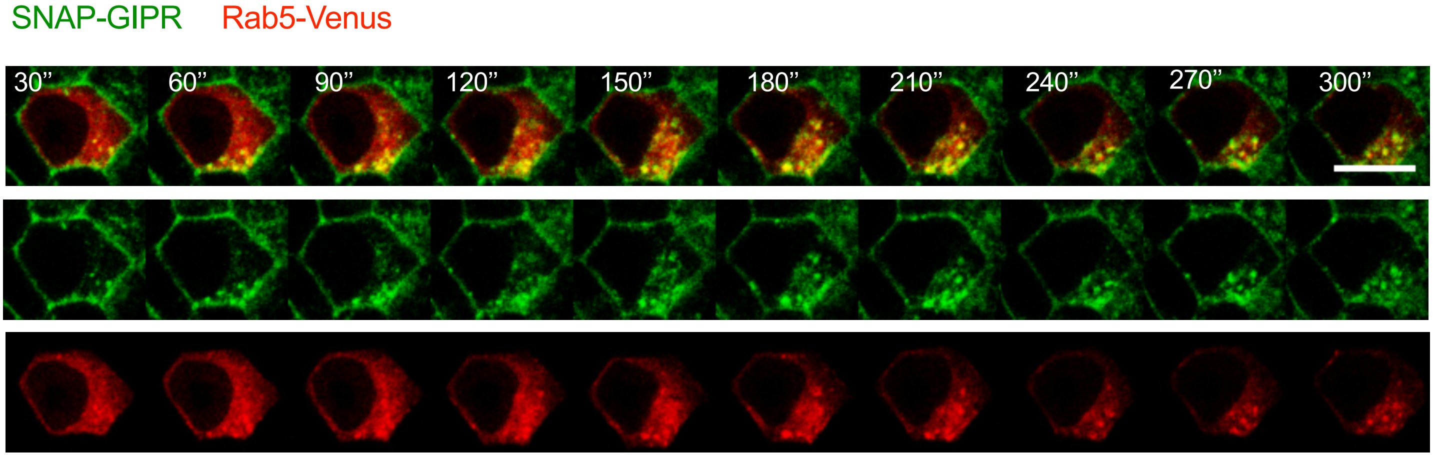

### Supplementary Figure 5

#### Supplementary Figure 5

## A

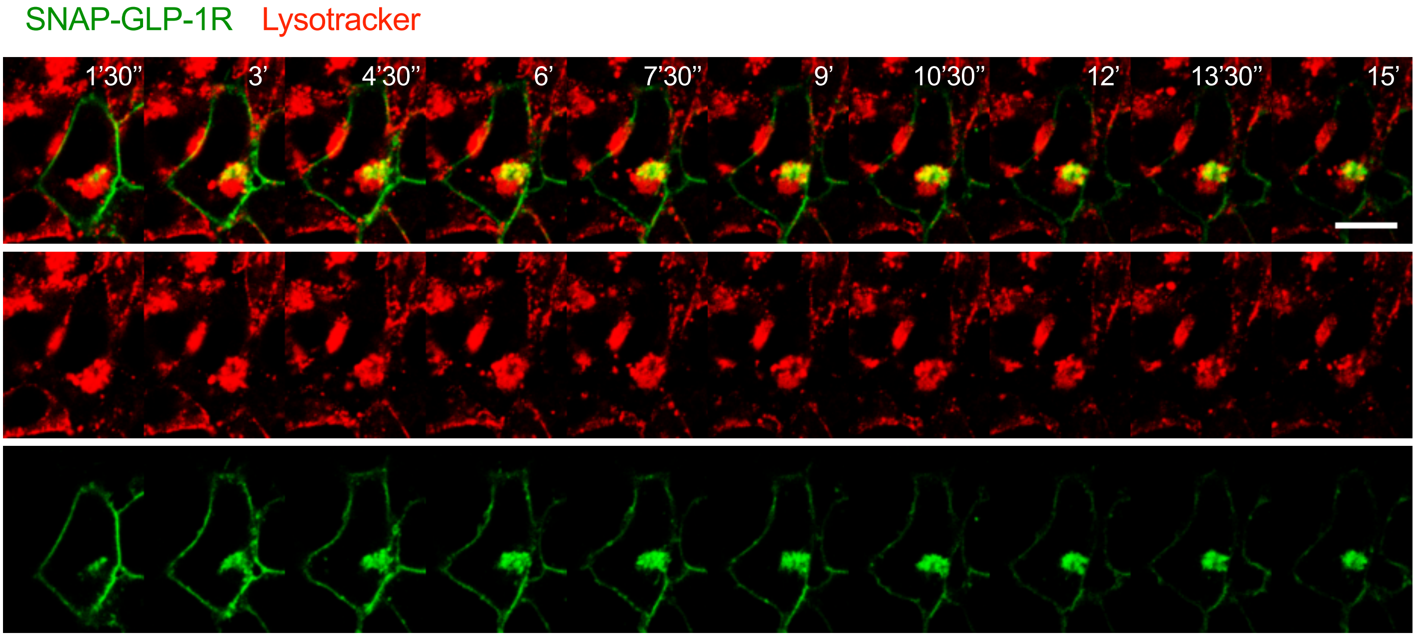


## B

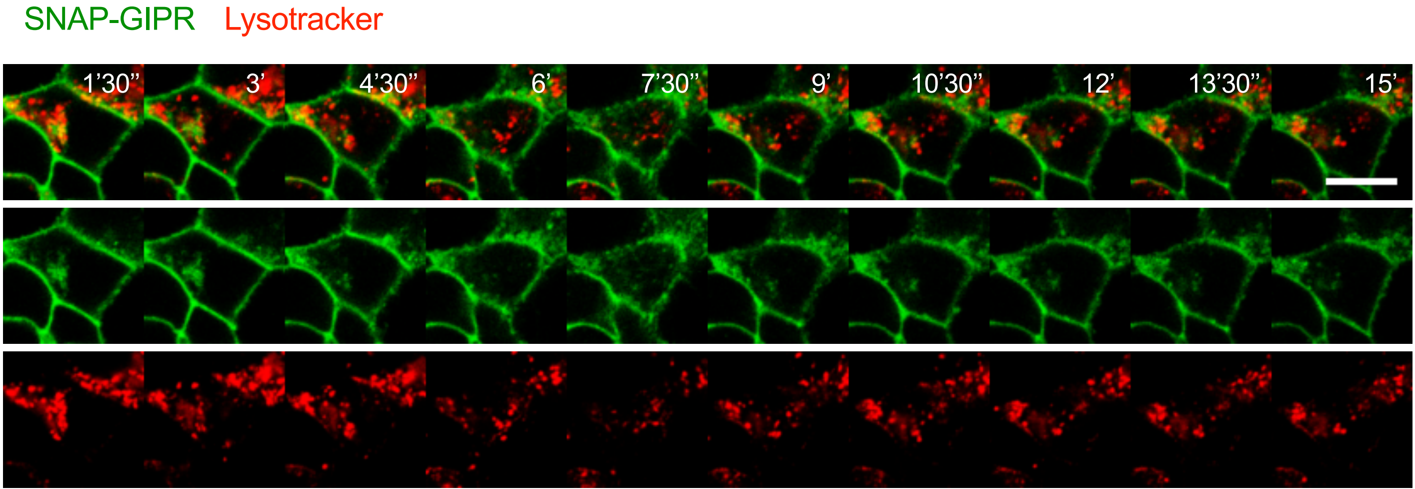

### Supplementary Figure 6

#### Supplementary Figure 6

**A**

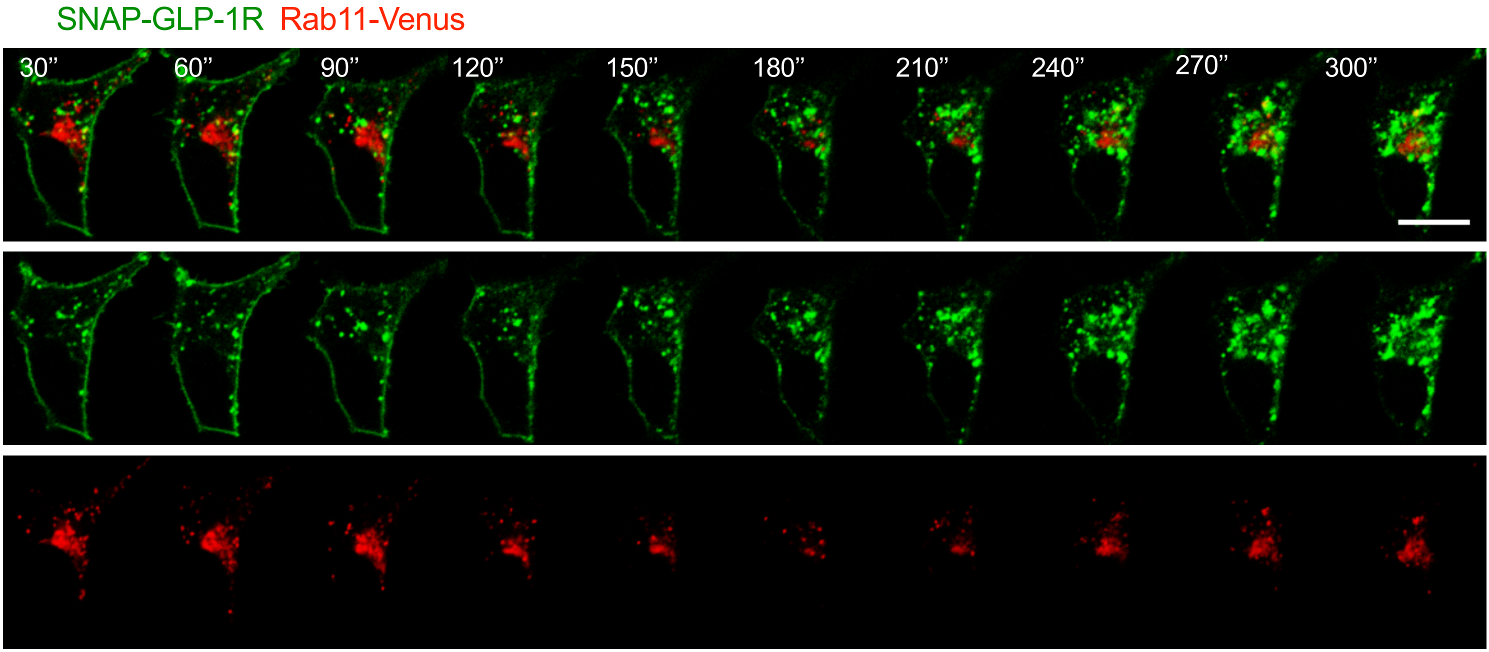


**B**

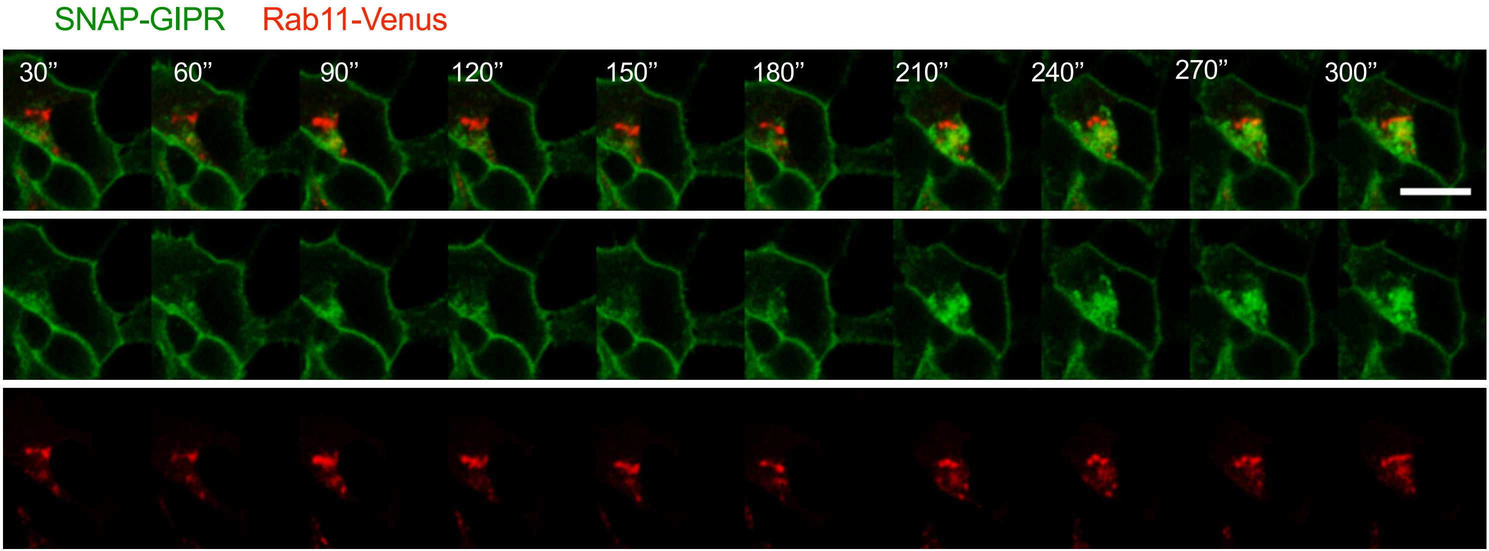

### Supplementary Figure 7

#### Supplementary Figure 7

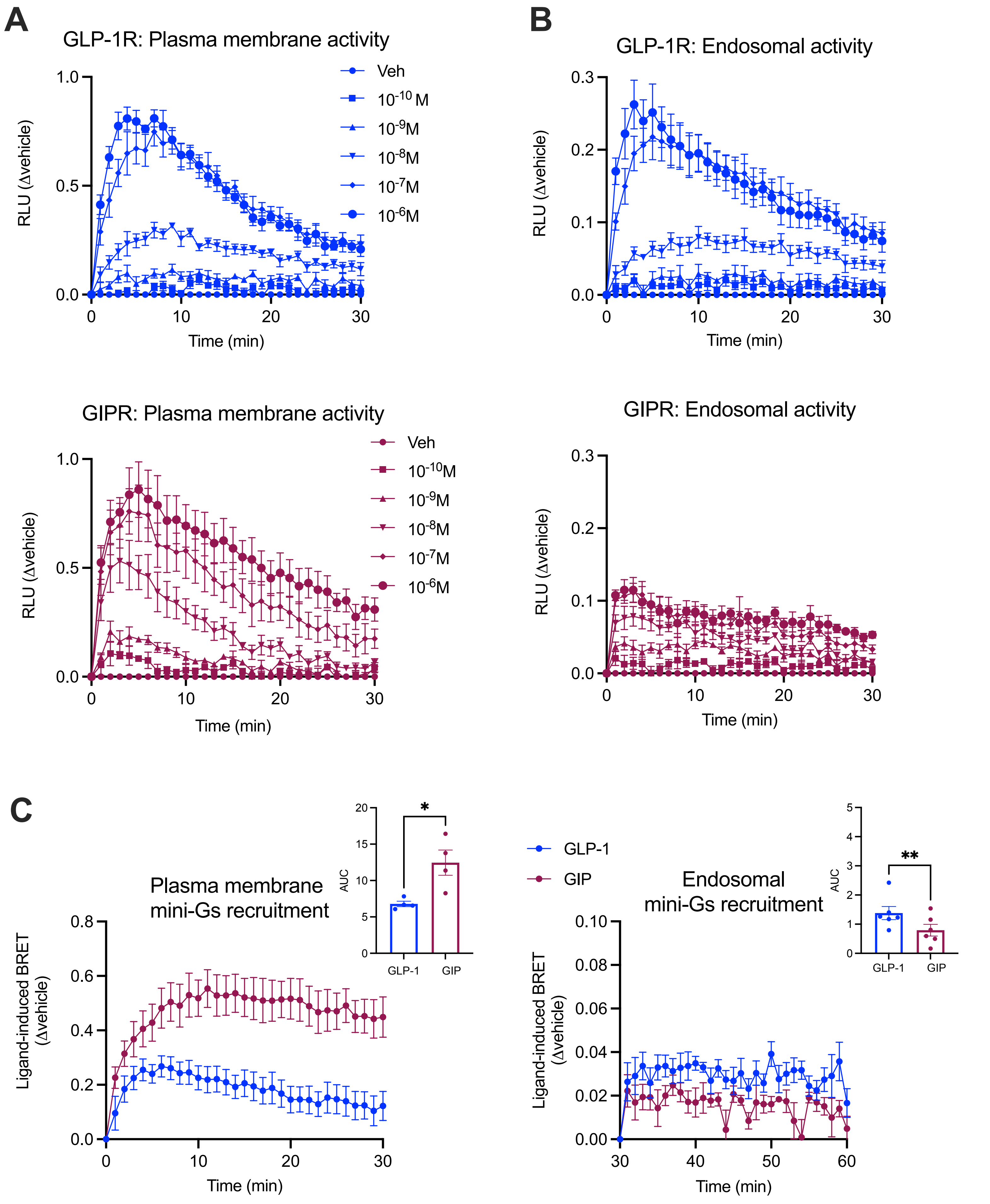
